# Structural Basis of How MGME1 Processes DNA 5′ Ends to Maintain Mitochondrial Genome Integrity

**DOI:** 10.1101/2024.01.15.575787

**Authors:** Eric Y.C. Mao, Han-Yi Yen, Chyuan-Chuan Wu

**Affiliations:** Department of Chemistry, College of Science, National Cheng Kung University, Tainan City 701, Taiwan; Department of Biochemistry and Molecular Biology, College of Medicine, National Cheng Kung University, Tainan City 701, Taiwan

**Keywords:** mtDNA replication, mtDNA degradation, MGME1, crystal structure

## Abstract

Mitochondrial genome maintenance exonuclease 1 (MGME1) helps to ensure mitochondrial DNA (mtDNA) integrity by serving as an ancillary 5′ exonuclease for DNA polymerase γ. Curiously, MGME1 exhibits unique bidirectionality *in vitro*, being capable of degrading DNA from either the 5′ or 3′ end. The structural basis of this bidirectionally and, particularly, how it processes DNA from the 5′ end to assist in mtDNA maintenance remains unclear. Here, we present a crystal structure of human MGME1 in complex with a 5′-overhang DNA, revealing that MGME1 functions as a rigid DNA clamp equipped with a single-strand-(ss)-selective arch, allowing it to slide on single-stranded DNA in either the 5′-to-3′ or 3′-to-5′ direction. Using a nuclease activity assay, we have dissected the structural basis of MGME1-derived DNA cleavage patterns in which the arch serves as a ruler to determine the cleavage site. We also reveal that MGME1 displays partial DNA-unwinding ability that helps it to better resolve 5′-DNA flaps, providing insights into MGME1-mediated 5′-end processing of nascent mtDNA. Our study builds on previously solved MGME1-DNA complex structures, finally providing the comprehensive functional mechanism of this bidirectional, ss-specific exonuclease.

## INTRODUCTION

Mitochondrial DNA (mtDNA) supports local production of electron transfer chain proteins in the mitochondrial matrix, thereby playing a vital role in sustaining mitochondrial function (1,2). Human mtDNA is a circular double-stranded DNA (dsDNA) of about 16.6 kilobase pairs (kbp). It is maintained by nucleus-encoded proteins that are trafficked to mitochondria. Consequently, genetic defects in these mtDNA-maintaining proteins could lead to impaired mtDNA integrity, prompting organelle dysfunction and disease (3,4).

First identified in 2013, mitochondrial genome maintenance exonuclease 1 (MGME1, also known as Ddk1) is a mitochondrial deoxyribonuclease in the PD–(D/E)XK nuclease superfamily that participates in mtDNA maintenance (5,6). Analyses of its nuclease activity *in vitro* revealed that MGME1 requires a free DNA end to exert its function (5,6), defining it as an exonuclease. MGME1 displays a strong preference for single-stranded DNA (ssDNA) over dsDNA. Intriguingly, unlike most exonucleases that typically work in one direction, MGME1 can digest DNA bidirectionally, enabling it to remove an ssDNA overhang or flap from various kinds of DNA duplexes (5-8). Loss-of-function mutations in the gene encoding MGME1 (*C20orf72*) cause systemic mtDNA maintenance disorders, characterized by mitochondrial dysfunction in multiple organs (5,9,10). Patients carrying such mutations harbor accumulations of mitochondrial 7S DNA and mtDNA depletion in their cells. Later studies revealed that MGME1 directly regulates the level of 7S DNA (5,6), which is generally viewed as a pre-termination replication product of mtDNA heavy (H)-strand synthesis (11). Despite its bidirectionality *in vitro*, depleting MGME1 was found to perturb 5′-end but not 3′-end processing of 7S DNA (12), indicating that MGME1 primarily serves as a 5′ exonuclease in this process. The 5′-end processing of 7S DNA and the nascent H-strand involves ribonuclease H1 (RNase H1)-mediated RNA primer removal and subsequent DNA trimming of ∼100 nucleotides (nt), from conserved sequence block 2 (CSB2) to the H-strand origin (OriH), before the DNA ends are joined by DNA ligase 3 to restore mtDNA integrity (1,2). Given that RNase H1 cannot fully remove all of the RNA primers and leaves a two-ribonucleotide (2-nt RNA) cap at the 5′-end of the nascent DNA (13-15), another nuclease is required to complete RNA primer removal. Many nucleases have been implicated in that process, including flap structure-specific endonuclease 1 (FEN1) (16,17), DNA replication helicase/nuclease 2 (DNA2) (18), and exonuclease G (EXOG) (19), but which enzyme plays the primary role remains under debate. Although MGME1 cannot digest RNA (5,6), an *in vitro*-coupled mtDNA replication-ligation assay revealed that it can generate ligatable ends by cooperating with the strand-displacement synthesis activity DNA polymerase γ (Polγ, the mtDNA replicase), which produces a 5′-DNA flap to facilitate MGME1’ action (7). By working together, the two enzymes are proposed to provide the ligatable ends for restoring the nascent DNA integrity during mtDNA replication (7). Apart from its role in mtDNA synthesis, further studies have uncovered that Polγ acts in mtDNA degradation via its 3′-exonuclease (3′-exo) activity (20-22). Intriguingly, MGME1 has also been found to participate in degrading mtDNA (21). Either inactivating the 3′-exo activity of Polγ or knocking out MGME1 greatly reduced mtDNA degradation (21). This finding is in line with the notion that Polγ and MGME1 work coordinately, with the 5′-exo activity of MGME1 complementing the 3′-exo activity of Polγ, thereby allowing simultaneous degradation of both DNA strands from the ends of linearized mtDNA (21). In addition to that interactomics analyses have shown that MGME1 associates with Polγ (12,23), consequently, MGME1 is currently believed to operate as an ancillary factor that provides the 5′-exonuclease (5′-exo) activity to the mtDNA replicase.

Despite the fact that MGME1 appears to primarily serve as a 5′ exonuclease in maintaining mtDNA, how it can operate bidirectionally is of interest in terms of its enzymatic and structural characteristics. Intriguingly, using an RNA-DNA-RNA chimeric substrate, Szczesny *et al*. (6) observed that MGME1 could skip the flanking RNA regions and cleave within the DNA region. This observation indicates that MGME1 can load itself onto single-stranded oligonucleotides from either the 5′- or 3′-free end, and that it possesses the ability to slide along the ss region to cleave at its preferential sites. Previously published crystal structures of *apo*-, ssDNA-bound, and 3′-overhang DNA-bound MGME1 (8) have revealed that the enzyme possesses an arched structure that allows ssDNA to thread through it. This arch, named as helical arch as it comprises α2 and α3 of MGME1 (8), is aligned downstream (*i*.*e*., the 3′ side according to the bound ssDNA) of the catalytic site. Consequently, in the 3′-overhang DNA-bound MGME1 structure (8), the catalytic site is oriented toward the duplex region, indicating that the enzyme would introduce a strand break between the third and fourth nucleotide upstream of the single-stranded-double-stranded (ss-ds) junction. The authors of that structural study suggested that processing DNA in the opposite direction, *i*.*e*., approaching and binding ssDNA from the 5′ end, would require a significant conformational change in MGME1 (8).

Due to a lack of structural information, how MGME1 processes DNA in the 5′-to-3′ direction remains unclear. Here, we have solved the crystal structure of MGME1 bound with a 7-nt 5′-overhang DNA, revealing that the protein operates as a rigid ssDNA clamp. Without conformational changes, MGME1 can load itself from either the free 5′ or 3′ end of a single-stranded oligonucleotide and digest the substrate bidirectionally by sliding along it in either direction. Importantly, combined with an *in vitro* nuclease activity assay, we clarify the structural basis of the distinct DNA cleavage patterns arising from MGME1 working in different directions, with the helical arch serving as a ruler to determine the site for DNA cleavage. Furthermore, we show that MGME1 is capable of partially unwinding the downstream duplex of an ss-ds junction in a sequence-context-dependent manner, enabling it to better resolve a 5′-ssDNA flap. Thus, our study provides important insights into the working mechanism of MGME1 as it cooperates with Polγ to ensure mtDNA maintenance.

## MATERIALS AND METHODS

### Molecular cloning

The human *C20orf72* gene (base pairs 61 to 1032, encoding residues 21 to 344 which lacks the N-terminal mitochondrial-targeting sequence) was cloned into pSol vector (Lucigen) to generate the pSol-His_8_-SUMO-MGME1 plasmid, which encodes a recombinant human MGME1 protein N-terminally fused to His_8_-tagged SUMO followed by a tobacco etch virus (TEV) protease cutting site. The plasmid was used to generate N-terminal-truncated MGME1 (ΔN-MGME1; residues 95 to 344) by means of In-Fusion Cloning (Takara Bio). Constructs expressing the MGME1 variants used in this study were generated by site-directed mutagenesis using the pSol-His_8_-SUMO-MGME1 plasmid as template. The sequences of the cloning and mutagenesis primers are presented in Supplementary Table S2.

### Protein purification

Plasmids were transfected into *Escherichia coli* BL21(DE3)pLysS cells for protein expression. The cells were grown in lysogeny broth (LB) at 37 °C until the cell density reached an OD_600_ of 0.8. Expression of recombinant MGME1 was induced by adding 0.05% L-rhamnose. Proteins were expressed at 20 °C for 16 hr. The cells were then harvested and lysed in lysis buffer (50 mM sodium phosphate pH 8.0, 500 mM NaCl, 10% [v/v] glycerol, 1% Tween 20, 0.5 mM phenylmethylsulfonyl fluoride (PMSF), 10 mM imidazole and 5 mM β-mercaptoethanol (2-ME)) by sonication. The lysate was centrifuged at 16,000 rpm for 45 min to remove cell debris. The supernatant was filtered through a 0.22-μm filter and loaded onto a Histrap FF column (Cytiva) connected with a ÄKTA Go (Cytiva) fast-protein liquid chromatography system. The column was washed to baseline using A buffer (lysis buffer lacking Tween 20 and PMSF). Column-bound proteins were eluted using B buffer (A buffer containing 200 mM imidazole). TEV protease was then added into the eluted proteins and dialyzed against 1 L of TEV reaction buffer (20 mM Tris-HCl pH 8.0, 200 mM NaCl and 1 mM dithiothreitol) at 4 °C for 4 hr. The TEV-treated proteins were again purified through a Histrap FF column. The flow-through fraction from the column was collected, concentrated, injected onto HiLoad Superdex 200 pg columns (Cytiva), and separated against gel-filtration buffer (20 mM Tris-HCl pH 8.0, 200 mM NaCl, 5 mM 2-ME and 1 mM ethylenediaminetetraacetic acid). The fractions containing pure MGME1 were collected, concentrated, and stored at −80 °C for further use. ΔN-MGME1 was purified according to the same protocol as described above. The purified ΔN-MGME1 was stored in a low-ionic-strength buffer (gel-filtration buffer containing 100 mM NaCl) to facilitate protein crystallization screening.

### *In vitro* nuclease assay

For each reaction, 100 nM of the indicated DNA substrate was incubated with the indicated concentration of MGME1 in a reaction buffer containing 100 μg/mL BSA, 10 mM HEPES pH 7.4, 150 mM NaCl and 2.5 mM MgCl_2_ at 37 °C. For the time-course experiments, an aliquot of the reaction mixture was pipetted out at the indicated time points and quenched by mixing with an equal volume of 2X TBE/Urea sample buffer (Thermo Fisher Scientific) and heating at 65 °C for 20 min. To analyze DNA cleavage patterns, quenched reactions were separated by 20 % native TBE polyacrylamide gel electrophoresis. The results were imaged using an Amersham Typhoon 5 Biomolecular Imager (Cytiva), with a 473 nm laser and 525BP20 filter for FAM, a 635 nm laser and 670BP30 filter for Cy5, and a 532 nm laser and 570BP20 filter for Cy3. The band signals were quantified using Image J. Statistical significance (*P* values) was determined in GraphPad Prism version 9.5.1 for MacOS (24) by two-tailed paired *t* test, with ** indicating *P* < 0.01.

### Crystallization

ΔN-MGME1 (7 to 11.5 mg/mL) was mixed with 241 to 397 μM of a 7-nt 5′-overhang DNA and 10 mM CaCl_2_. Crystallization was conducted by the hanging-drop vapor diffusion method. The drops contained an equal volume of the protein-DNA solution and the reservoir solution. Crystallization screening was performed using mosquito crystal (STP Labtech). Crystals were grown in a reservoir solution containing 100 mM sodium citrate/citric acid pH 4.0, 200 mM sodium citrate tribasic and 20% [w/v] polyethylene glycol 3,350 within 2 to 7 days at 10 °C.

### Data collection, structure determination, and model analysis

For data collection, the grown crystals were transferred to a cryoprotective solution (the reservoir solution containing 25% [v/v] glycerol) and quickly frozen in liquid nitrogen. X-ray diffraction data was collected at TLS beamline 15A, National Synchrotron Radiation Research Center (NSRRC), Taiwan. The data was processed in the HKL2000 program suite (25). The crystal structure was solved by molecular replacement using the AlphaFold2-predicted human MGME1 structure (UniProt accession number: Q9BQP) (26,27) with Phaser (28) in the Phenix program suite (29). In this initial solution, we found some regions exhibited significant higher average B-factors, especially the helical arch of one of the two polypeptide chains. These regions with significant higher B-factor were deleted from the model, and we used the model as the template to search solutions with Phaser again. The second solution displayed a lower average B-factor than the initial one. The deleted regions and DNA were built manually according to the electron density maps using Coot (30). The model was subjected to cycles of manual building and refinement with Coot and phenix.refine (31). The crystallographic and model refinement parameters are listed in Supplementary Table S1. Structural superimpositions were performed by aligning protein secondary structures in PyMoL (32). All structural figures were generated using PyMoL.

### Electrophoretic mobility shift assay (EMSA)

For each reaction, 100 nM of the indicated DNA substrate was incubated with the indicated concentration of MGME1 in EMSA buffer (20 mM Tris-HCl pH 8.0, 10 mM CaCl_2_ and 150 mM NaCl) at 25 °C for 10 min. The reaction products were resolved by 5% TBE-native polyacrylamide gel electrophoresis. The results were imaged using an Amersham Typhoon 5 Biomolecular Imager (Cytiva) (473 nm laser and 525BP20 filter for FAM). The band signals were quantified using Image J. The data was fitted to specific binding activity with a Hill slope model by using GraphPad Prism version 9.5.1 for MacOS (24).

## RESULTS

### Crystallization of 5′-overhang DNA-bound ΔN-MGME1

To capture MGME1 processing ssDNA in the 5′-to-3′ direction, we first designed a 5′-overhang DNA for co-crystallization with the enzyme. To do so, a 25-nt 3′-6-carboxyfluorescein (FAM)-labeled oligonucleotide probe was paired with complementary strands of varying length to generate 5′-overhang DNA ranging from 5 to 9 nt (Supplementary Figure S1A). The 5′ end of the probe was phosphorylated (denoted 5′P) to enhance the efficiency of MGME1 in processing the substrate (33). An *in vitro* nuclease activity assay revealed that MGME1 favored the 7-nt overhang DNA, as evidenced by efficient and homogeneous cleavage of the overhang DNA by the enzyme that directly removed 3 nt from its 5′ end (Supplementary Figure S1B). MGME1 exhibited poor cleavage performance when the overhang was shorter than 6 nt, in agreement with a previous report on MGME1 processing of a 5′-ssDNA flap (7). Accordingly, we used a DNA duplex carrying a 7-nt 5′ overhang for co-crystallization with MGME1. However, we failed to obtain crystals, despite an exhaustive crystallization screening. By examining available structures of MGME1, we observed that the N-terminus of the enzyme spanning residues 21 to 94 (*i*.*e*., excluding its mitochondrial-targeting sequence - MTS residues 1-20; Figure 1A) had neither been visualized in previous crystal structures (8) nor been predicted by AlphaFold2 to form a high-confidence structure (26,27), suggesting a dynamic nature of the particular region. We reasoned that the N-terminus may have inhibited MGME1 crystallization under our experimental conditions, so we deleted the N-terminus to produce recombinant ΔN-MGME1 (residues 95 to 344) for crystallization (Figure 1A, Supplementary Figure S1 C-E). ΔN-MGME1 exhibited significantly stronger nuclease activity than the full-length enzyme (residues 21 to 344, *i*.*e*., lacking only the MTS) (Supplementary Figure S1F), implying the N-terminus exerts an autoinhibitory effect, as postulated previously (8). By supplying our co-crystallization mixture with the inhibitory ion Ca^2+^, we successfully blocked ΔN-MGME1 activity (Supplementary Figure S1G) and obtained crystals capturing the substrate-bound ΔN-MGME1 in the pre-cleavage state (Figure 1B and Supplementary Table S1).

**Figure 1.**
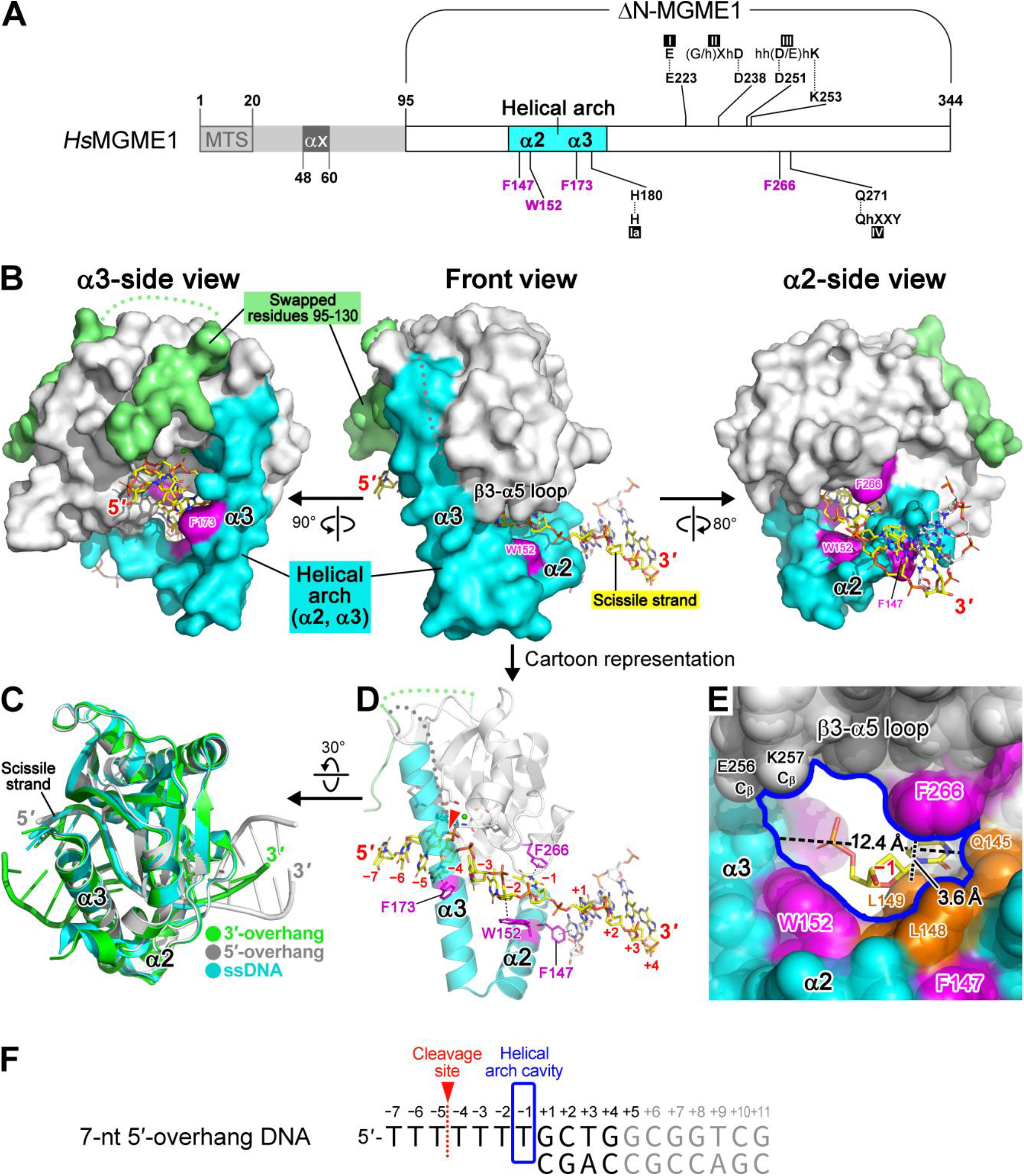
Crystal structure of MGME1 in complex with 5′-overhang DNA. (**A**) The domain organization of *Homo sapiens* (*Hs*) MGME1. MTS, mitochondrial-targeting sequence. The catalytic residues of conserved motifs (Ia, I, II, III and IV) in the PD–(D/E)XK nuclease superfamily are labeled. The DNA-stacking aromatic residues are highlighted in magenta. (**B**) Overall structure of the ΔN-MGME1-5′-overhang DNA complex. The protein is shown in surface representation, with DNA in stick representation. The non-scissile strand is shown in gray. (**C**) Superimposition of MGME1-DNA complexes. The 5′-overhang DNA-bound MGME1 (this study) was aligned with the 3′-overhang DNA-bound (PDB ID: 5ZYT) and ssDNA-bound (PDB ID: 5ZYU) MGME1 structures, yielding root-mean-square-deviation (RMSD) values of 0.409 (over 181 C_α_) and 0.359 (over 192 C_α_), respectively. (**D**) Cartoon representation of the front view of the ΔN-MGME1-5′-overhang DNA complex. (**E**) The cavity under the helical arch. To better represent the protein surface landscape, the protein is shown as spheres (van der Waals radius) covered with a transparent surface (reflecting solvent contact). The edge of the cavity is highlighted by a blue line. For clarity, only the nt −1 is shown for the bound DNA. Residues contributing to cavity formation are colored orange, except for F266 and W152 that are presented in magenta. Note that the side chains of E256 and K257 are not modeled due to lack of respective electron density maps. (**F**) The 7-nt 5′-overhang DNA used for crystallization. The distal region of the downstream duplex that could not be modeled due to ambiguous electron density maps is colored gray.

The determined crystal structure contains two essentially identical ΔN-MGME1-DNA complexes in an asymmetric unit, with the two complexes stacking with each other in a two-fold symmetric manner, as resolved previously for the MGME1-ssDNA complex structure (PDB ID: 5ZYU) (8) (Supplementary Figure S2A). In this particular stacking of ΔN-MGME1-DNA complexes, residues 95 to 130 of the enzyme protrudes toward the other protein protomer and packs on the protein surface (Supplementary Figure S2B). Curiously, this same packing between residues 95-130 and the protein surface can be found intramolecularly in one of the *apo*-MGME1 crystal structures (Mn^2+^-bound; PDB ID: 5ZYW) and in the AlphaFold2-predicted MGME1 model (the bottom panel in Supplementary Figure S2B). Given that MGME1 has been purified as a monomeric protein by others (6,8), as well as herein by us (Supplementary Figure S1 C-E), we conclude that *in crystallo* swapping of residues 95-130 between the adjacent ΔN-MGME1-DNA complexes arose during crystallization and that, in solution, residues 95-130 should pack intramolecularly against the protein surface.

### MGME1 operates as a rigid ssDNA clamp capable of degrading DNA from either end

Of the two ΔN-MGME1-DNA complexes in our crystal structure, one presented significantly better electron density maps and a lower average B-factor (Supplementary Figures S2C and S3). Therefore, we used this complex (protomer B in Supplementary Figure S2) for all subsequent analyses and figure illustrations. The previous structural study of MGME1 suggested that a significant protein conformational change may be required for the enzyme to approach substrate from the 5′ end (8). However, our solved structure of the 5′-overhang DNA-bound complex revealed no notable structural changes in the enzyme to accommodate either the 5′-ssDNA overhang of the substrate or its duplex region (Figure 1C). Neither structural change in the ssDNA overhang, ranging from nt –7 to –1 of the scissile-DNA strand, were observed (Figure 1C and D). In our structure, the 5′-ssDNA overhang threads through the helical arch from the α2 side (Figure 1B), with nt –1 being confined by a cavity formed between α2 and the β3-α5 loop; more specifically, between residues W152, Q145, L148, L149 and F266. This cavity, which is about 12.4 Å long and 3.6 Å wide, is the narrowest region of the space under the helical arch (Figure 1E), accommodating only single-stranded oligonucleotides to pass through it. As a result, in our solved 5′-overhang DNA-bound structure, the duplex region was anchored on the α2 side of the helical arch (Figure 1B) with the first base pair, labeled +1 bp in Figure 1D and F, packed against the surface of α2. In contrast, in the 3′-overhang DNA-bound structure reported previously (8), the duplex region was anchored alongside this cavity from the α3 side. Consequently, our structure reveals that MGME1 operates as a rigid ssDNA clamp that can digest ssDNA in either direction without changing protein conformation.

In line with the insights from the structures, MGME1 has consistently been observed as ceasing ssDNA digestion near the ss-ds junction (Supplementary Figure S1 and Figure 2), *i*.*e*., where the helical arch likely acts as a structural barrier for duplex DNA. Intriguingly, since the arch is located downstream of the catalytic site, it precedes the catalytic site and encounters the duplex region first when MGME1 digests DNA in the 5′-to-3′ direction. Accordingly, MGME1 stops digesting DNA before the catalytic site reaches the ss-ds junction, so that its catalytic products always have a short 5′ overhang (Figure 2A). In contrast, when MGME1 digests DNA in the 3′-to-5′ direction, the catalytic site moves in front of the helical arch and is anchored in the duplex region when the enzyme is halted, resulting in cleavages in the duplex region that also produce 5′-overhang DNA as a final product (Figure 2B).

**Figure 2.**
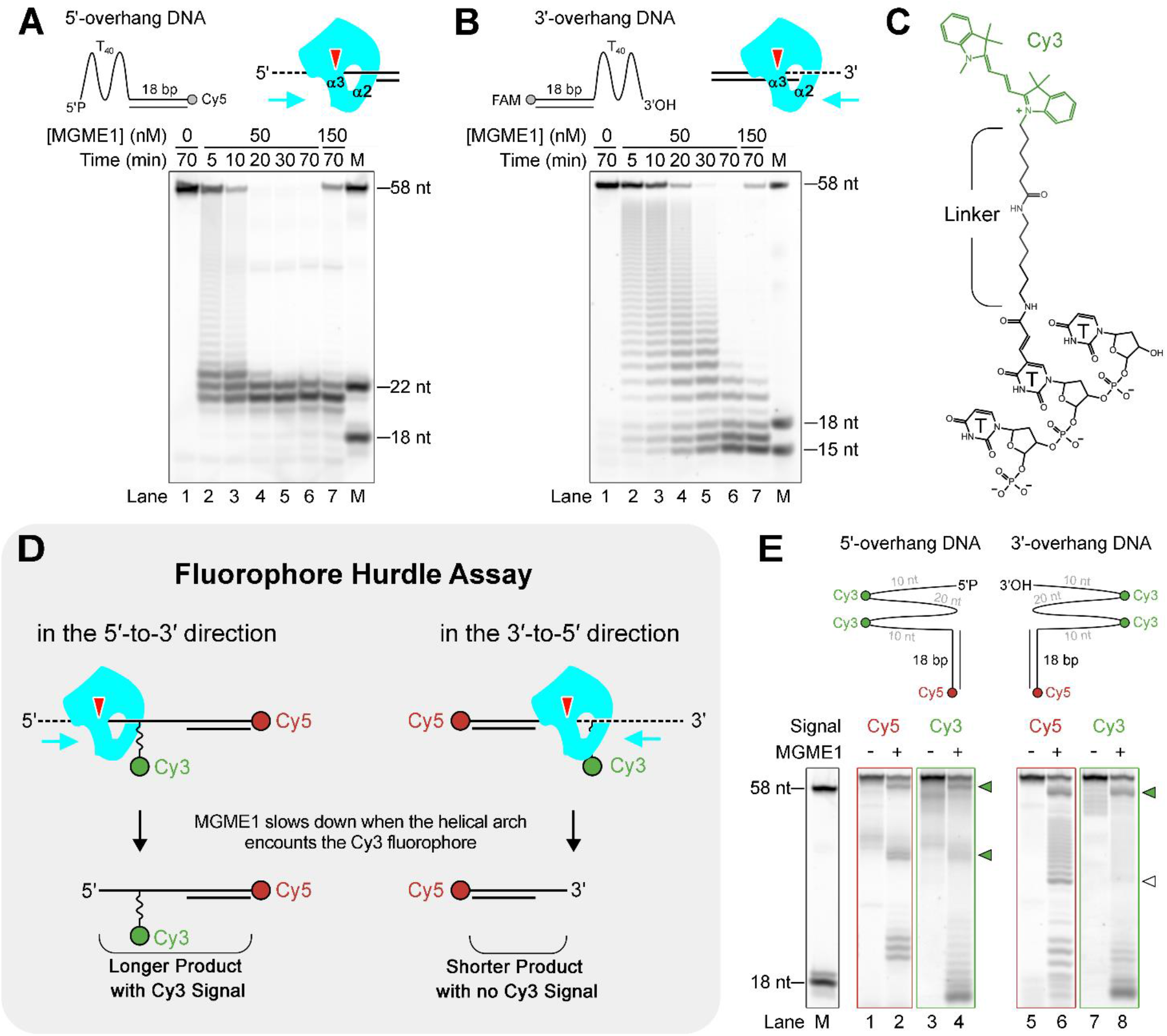
MGME1 exhibits distinct DNA digestion patterns when acting in different directions. (**A** and **B**) MGME1-derived DNA cleavage patterns from 5′-overhang (**A**) and 3′-overhang (**B**) DNA substrates. A polyT sequence (T_40_) was employed as the 40-nt ssDNA region of substrates to avoid secondary structure formation. Red arrowheads indicate the MGME1-mediated cleavage sites. (**C**) The chemical structure of the Cy3-photophore to label the thymine base. (**D** and **E**) Schematic (**D**) and results (**E**) of the fluorophore hurdle assay. The T_40_-ssDNA overhang region of the substrates was labeled with two Cy3 fluorophores separated by 20 nt. For each reaction, 100 nM of the substrate was incubated with 50 nM MGME1 for 20 min. Green arrowheads indicate accumulations of MGME1-derived cleavage intermediates, and the hollow arrowhead indicates intermediates without Cy3 signal. M: synthetic nucleotide markers.

To provide evidence for this scenario, we employed oligonucleotide probes carrying internal Cy3 labels in their ssDNA overhang regions (Figure 2C), assuming that the bulky Cy3 fluorophore would act as a hurdle to MGME1 upon the helical arch encountering it (Figure 2D). Indeed, we observed two clusters of MGME1-derived cleavage intermediates upon gel electrophoresis of the probes labeled at the distal end with a Cy5 fluorophore, corresponding to the two Cy3-labeled sites (indicated by green arrowheads in Figure 2E). Importantly, when MGME1 digested DNA from the 5′ end, we observed a second cluster of cleavage intermediates by detecting Cy3 signal, indicating that MGME1 had stopped digesting the DNA before the catalytic site had reached the second Cy3-labeled site so that the fluorophore remained on the probe. In contrast, we did not observe Cy3 signal for the second cluster of cleavage intermediates when MGME1 digested DNA from the 3′ end (indicated by the open arrowhead in Figure 2E), meaning that MGME1 had completely removed the Cy3 fluorophores from the probe. This latter observation supports the notion that the catalytic site travels in front of the helical arch as MGME1 slides along the ssDNA in the 3′-to-5′ direction.

Collectively, these observations indicate that the helical arch serves as a structural barrier restricting MGME1 to ssDNA degradation. Moreover, this structural feature also defines MGME1 as a single-strand-specific exonuclease as it allows only ss oligonucleotides to pass through. In our *in vitro* nuclease activity assay, digestion of a blunt-end dsDNA was only observed when we used a high concentration of MGME1 (400 nM, Supplementary Figure S1H).

### MGME1 displays partial DNA unwinding ability to facilitate 5′-ssDNA flap removal

MGME1 digested the 7-nt 5′-overhang and 5′-flap dsDNA with comparable efficiency, with no notable differences in the derived DNA cleavage patterns (Supplementary Figure S1I). Therefore, we assumed that the enzyme interacts similarly with these two types of substrate. In the absence of a resolved 5′-flap DNA-bound MGME1 structure, we postulated that MGME1 interacts with this substrate in a manner similar to FEN1 (34,35), in which the non-scissile strand is kinked at the ss-ds junction and the two parts of the duplex are positioned almost vertically to form a splayed-arm architecture (Supplementary Figure S4A). Accordingly, when MGME1 interacts with a 5′-flap DNA substrate, we predicted that the upstream region of its splayed arm may be anchored on the surface of the helical arch without the need for the enzyme to adopt a significant protein conformational change (Supplementary Figure S4B).

Although MGME1 has been implicated in processing the 5′-flap of the nascent mtDNA H-strand, MGME1 could not generate the ligatable end without the 3′-exo activity Polγ in an *in vitro* coupled mtDNA replication-ligation assay (7), indicating that MGME1 cannot directly resolve the flap by cleaving at the ss-ds junction. Consistent with this notion, our 5′-overhang DNA-bound MGME1 structure reveals that four nucleotides (ranging from nt −4 to −1) are flanked by the catalytic site-pointing scissile bond and the helical arch cavity (Figure 1D-F). Consequently, MGME1 would obligatorily leave an unresolved 4-nt 5′-overhang on the product, therefore, further trimming of either the 5′-flap (by MGME1) or the 3′-terminus (by Polγ) is required for resolving the flap and generating a ligatable nick for DNA ligase (7). However, although MGME1 indeed mostly halts ssDNA digestion around the last 4 nucleotides of the 5′-overhang DNA (see the cluster of products of ∼22 nt, lanes 2 to 7 in Figure 2A) or the 5′-flap DNA (lanes 2 to 5 in Figure 3A), we observed that the position of the final cutting site essentially depends on the sequence context of the downstream duplex. If the first base pair of the downstream duplex is an AT pair, MGME1 can proceed 1 nucleotide further, as evidenced by the rapid generation of 21-nt but not 22-nt cleavage products from the 5′-overhang DNA (lane 2 in Figure 2A) or 5′-flap DNA (lane 2 in Figure 3A) substrates, both of which host an 18-bp duplex region. Neither a higher enzyme concentration (150 nM, lane 7 in Figure 2A) nor a prolonged incubation time (70 min, lane 5 in Figure 3A) prompted further digestion into the duplex region. Interestingly, by substituting five base pairs downstream of the ss-ds junction with AT pairs, we could make MGME1 mostly cut at the ss-ds junction of the 5′-flap DNA substrate (lanes 2 to 5 in Figure 3B). This outcome indicates that MGME1 is able to partially unwind the downstream duplex by up to four base pairs to overcome the gap between the catalytic site and the helical arch to resolve the 5′ flap, where an AT-rich downstream duplex is a prerequisite. MGME1 retained this DNA sequence context-dependent behavior when catalyzing a short (7-nt) flap (Figure 3C and D), which better represents its putatively endogenous 5′-flap substrate generated by the strand-displacement synthesis activity of Polγ (7,36).

**Figure 3.**
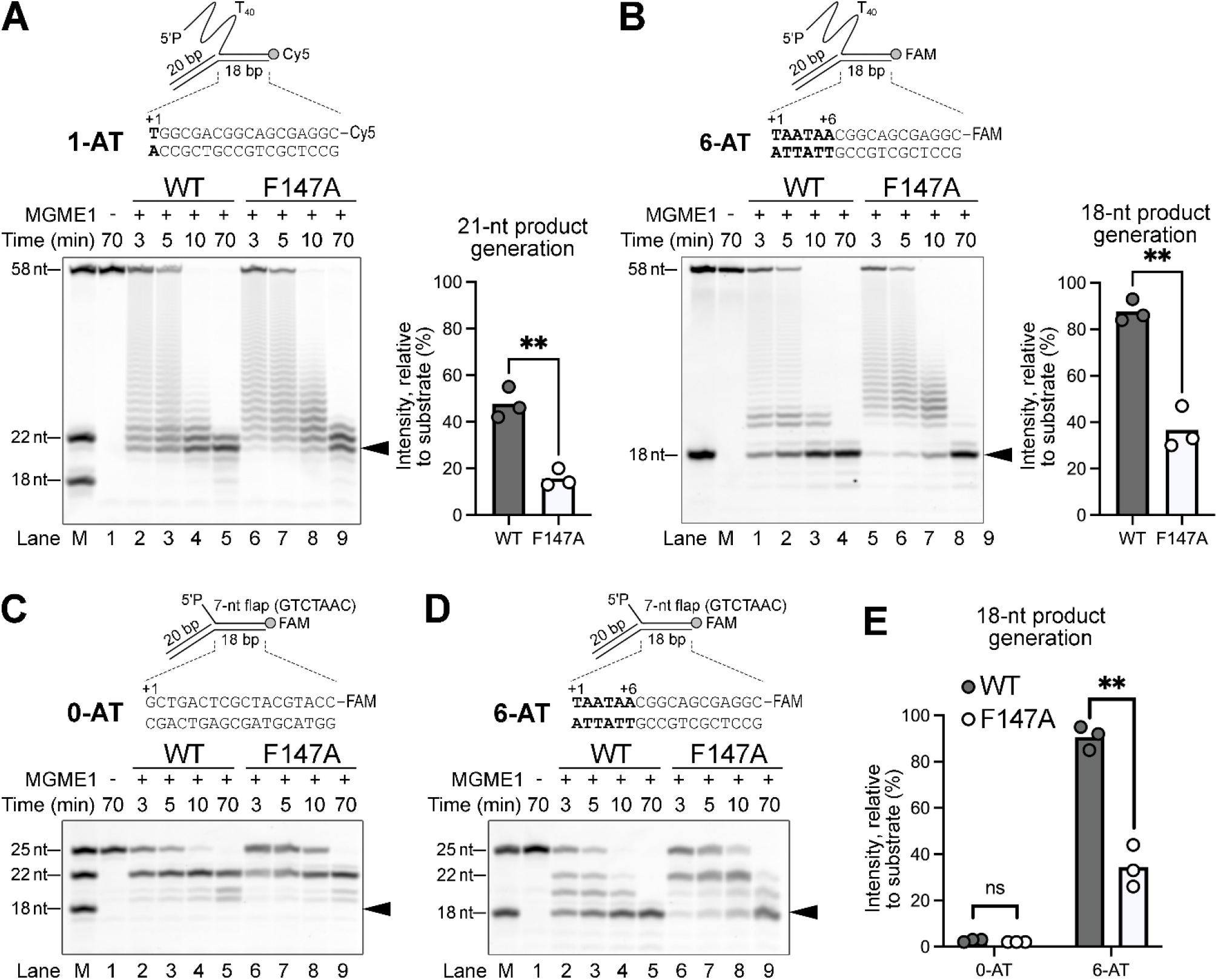
MGME1 unwinds the duplex downstream of the ss-ds junction in a GC content-dependent manner. (**A** and **B**) Activity of MGME1 and MGME1-F147A in digesting 40-nt 5′-flap DNA having one (1-AT, **A**) or six continuous (6-AT, **B**) AT pairs in the downstream duplex region. (**C** and **D**) Activity of MGME1 and MGME1-F147A in digesting 7-nt 5′-flap DNA with no (0-AT, **C**) or 6-AT (**D**) pairs in the downstream duplex region. Each reaction contains 100 nM substrate and 50 nM MGME1. (**E**) Quantification of the MGME1-derived 18-nt product band intensity, as shown in panels **C** and **D**. For all panels, black arrowheads indicate the product bands that were quantified in the corresponding bar charts. Two-tailed paired t-test was used to compare the mean values from two populations, **: *p* ≤ 0.01; ns: statistically non-significant.

To provide additional structural insights to support MGME1’s ability to partially unwind the duplex DNA downstream of the ss-ds junction, we inspected our resolved ΔN-MGME1-DNA structure and observed that residue F147, which protrudes from helix α2, points toward the first base pair of the downstream duplex, where it stacks with the complementary base of nt +1 (Figure 1D). We speculated that this residue may play a role in the partial unwinding of the downstream duplex, so we substituted it with alanine and observed that the F147A variant was significantly weaker at resolving the 5′ flap, as indicated by less final cleavage product being generated by this variant over a given time period (Figure 3, Supplementary Figure S5). Notably, the F147A variant exhibited comparable efficiency in processing the ssDNA overhang as wild-type enzyme (Supplementary Figure S5B and D). This outcome supports that residue F147 primarily contributes to processing of the downstream duplex at the ss-ds junction rather than digesting the ssDNA overhang.

Taken together, our structural analysis and nuclease activity data support that MGME1: i) is prone to cleaves upstream of a ss-ds junction due to the gap between its catalytic site and the ss-selective arch; but ii) possesses the ability to partially unwind the downstream duplex to allow cleavage at the ss-ds junction in a sequence-context-dependent manner; and iii) this latter capability requires a ssDNA region in front of the duplex is unwound, as evidenced by the enzyme’s extremely poor cleavage activity on blunt-end duplex DNA. This last observation is supported by structural insights from our resolved ΔN-MGME1-DNA complex structure, where ssDNA being threaded through the helical arch is a prerequisite for proper substrate alignment in the enzyme’s catalytic site.

### Identifying the DNA-stacking aromatic residues responsible for positioning ssDNA in the helical arch

Revealing by our and the previously solved MGME1-DNA complex structures (8), the nuclease possesses several aromatic residues that stack either with bases (including F147, F173 and F266) or the sugar pucker (W152) of the bound DNA (Figure 1D). By substituting these residues with alanine and assaying the exonuclease activity of the respective variants, we determined that removing the aromatic ring of F173 and F266 specifically enhanced the 5′-exo but not the 3′-exo activity of MGME1 (Figure 4A and B). Curiously, double substitution of these two residues to alanines did not elicit a synergistic effect but instead reverted the 5′-exo activity to the original level (Figure 4C). Given that both residues interact simultaneously with the substrate (7-nt 5′-overhang DNA) and the cleavage product, which is a 4-nt 5′-overhang DNA according to both the results of our nuclease assay (Supplementary Figure S1B) and the resolved ΔN-MGME1-DNA complex structure (Figure 1F), we reasoned that the enhanced 5′-exo activity of the mutant variant may involve more rapid product release contributing to a higher reaction turnover rate. To test this notion, we performed an electrophoresis mobility shift assay (EMSA) to investigate the ability of the F173A, F266A and F173A/F266A mutant variants to bind substrate (7-nt 5′-overhang DNA, Figure 4E) and cleaved product (4-nt 5′-overhang DNA, Figure 4F), respectively. The cleaved product represents the 22-nt product band generated by MGME1 upon cleaving the 7-nt 5′-overhang DNA, as shown in Supplementary Figure S1B and Figure 4A and C. By quantifying the reduction in signal intensity for the bands representing free DNA, which is complementary to the emergence of MGME1-DNA complex 1 when interacting with the 7-nt 5′-overhang DNA (Figure 4G), we observed that although both the F173A and F266A variants retained the ability to bind to the substrate, they exhibited significantly impaired interaction with the cleaved product (Figure 4 E, F, H and I, and Table 1). These findings support the notion that the enhanced 5′-exo activity is mainly attributable to faster product release from these variants. In contrast, double substitution of F173 and F266 with alanines reduced interaction with both substrate and product, with diminished substrate binding thus offsetting the benefits of faster product release. We also noticed that the F173A/F266A variant displayed poor 3′-exo activity, yet single substitution of either F173 or F266 with alanine did not have a notable impact on 3′-exo activity (Figure 4 B and D). This outcome could be explained by weakening of the interaction between the F173A/F266A variant and the 3′-overhang substrate owing to simultaneous loss of two DNA positioning residues. Collectively, these data show that residues F173 and F266, which respectively stack with nt −4 and −1 (Figure 1D), contribute to proper positioning of the ssDNA that threads through the helical arch.

**Table 1:**
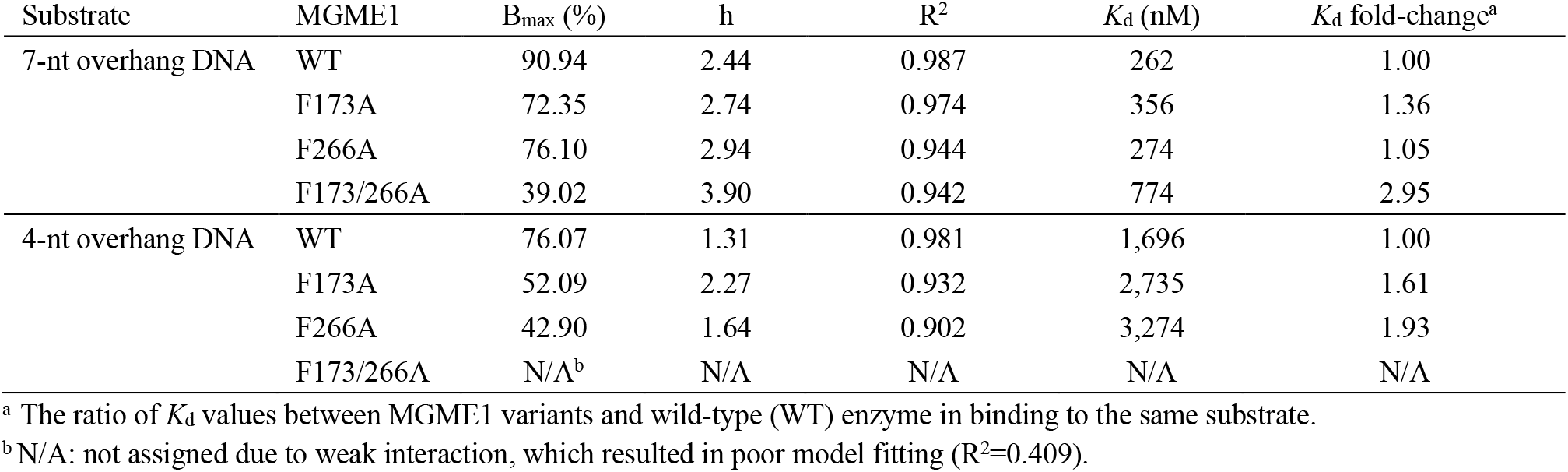
Binding affinity (*K*_d_) between MGME1 and 7-nt and 4-nt overhang DNA, as determined by electrophoretic mobility shift assay (EMSA)

**Figure 4.**
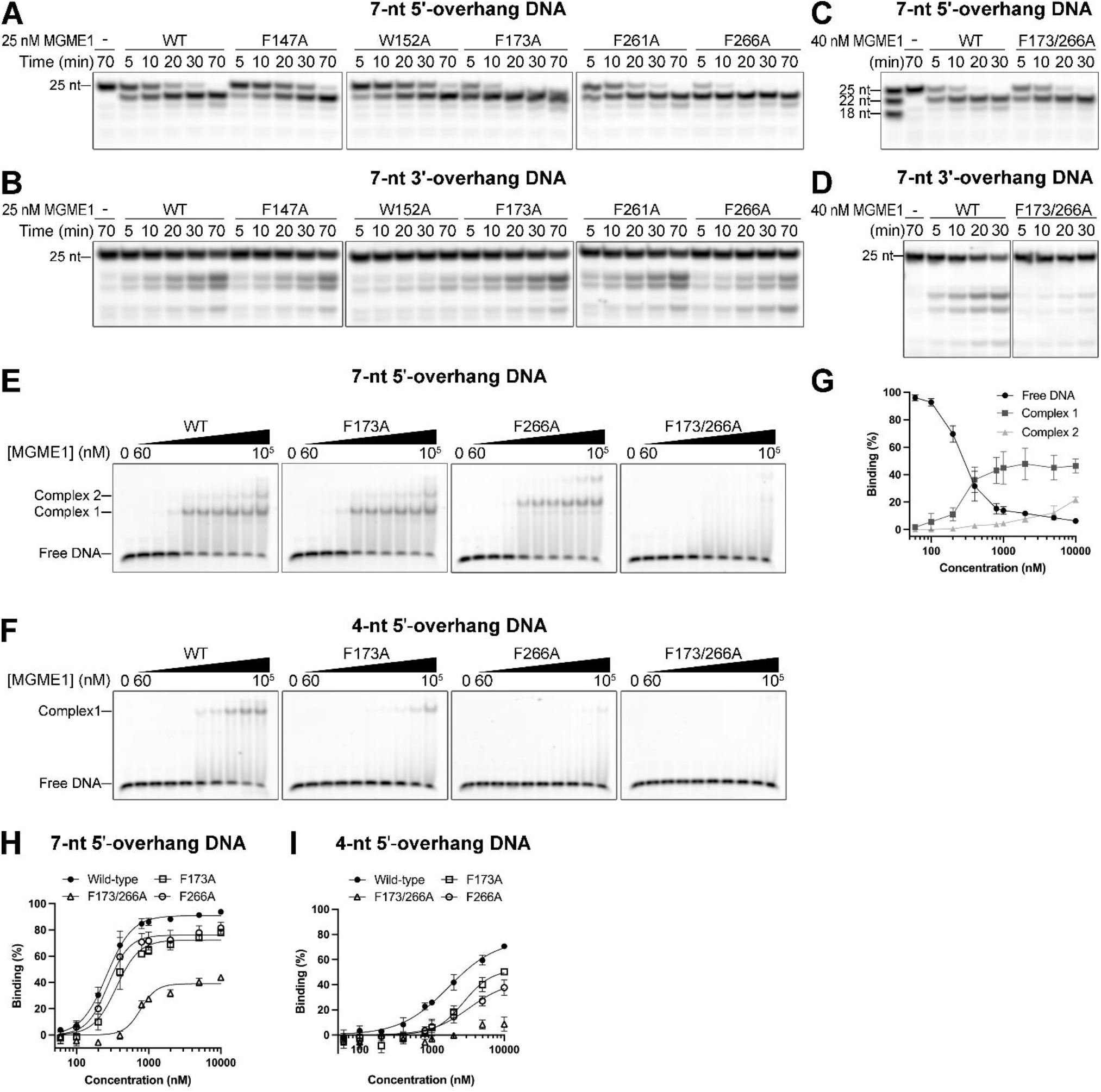
F173A and F266A substitutions exert distinct effects on 5′-exo and 3′-exo activities. (**A** and **B**) Nuclease activity assay to assess the 5′-exo (**A**) and 3′-exo (**B**) activities of MGME1 variants carrying a substitution of a DNA-stacking aromatic residue. (**C** and **D**) The 5′-exo (**C**) and 3′-exo (**D**) activities of the F173/266A variant. (**E** and **F**) Electrophoresis mobility shift assay (EMSA) to determine the ability of MGME1 variants to bind 7-nt (**E**) and 4-nt (**F**) 5′-overhang DNA. (**G**) Quantification of the EMSA results from WT MGME1 interacting with the 7-nt 5′-overhang DNA, as shown in panel **E**. (**H** and **I**) Quantification of the EMSA results from panels **E** and **F**, respectively. The data (n=3) were fitted to specific binding activity with a Hill slope model using GraphPad Prism. The best-fit values are listed in Table 1.

## DISCUSSION

MGME1 participates in mtDNA maintenance by serving as an ancillary 5′ exonuclease to the mtDNA replicase Polγ. Through its tight functional association with Polγ, MGME1 plays a key role in ensuring mtDNA integrity. MGME1 deficiency prompts mtDNA depletion, deletions, and rearrangements (5,12,23,37). Notably, it may require long-term MGME1 deficiency to impact mtDNA copy number in cells, since short-term (days) depletion or overexpression of MGME1 in HeLa cells did not alter mtDNA content (5,6,38). Endeavoring to establish how MGME1 processes mtDNA, we have solved the crystal structure of MGME1 in complex with a 5′-overhang DNA. Our structure reveals that the enzyme operates as a rigid DNA clamp capable of sliding along single-stranded oligonucleotides in either direction without necessitating a conformational change in the enzyme core. Consequently, MGME1 is promiscuous in cleaving ssDNA from various kinds of DNA duplexes, including overhang, flap, and forked duplexes (Figure 5A). Given this promiscuity, MGME1 levels in cells likely require strict regulation. Overexpressing MGME1 in patient-derived fibroblasts prompted severe mtDNA depletion (5), implying adverse non-specific ssDNA degradation can be mediated by an excess of MGME1.

**Figure 5.**
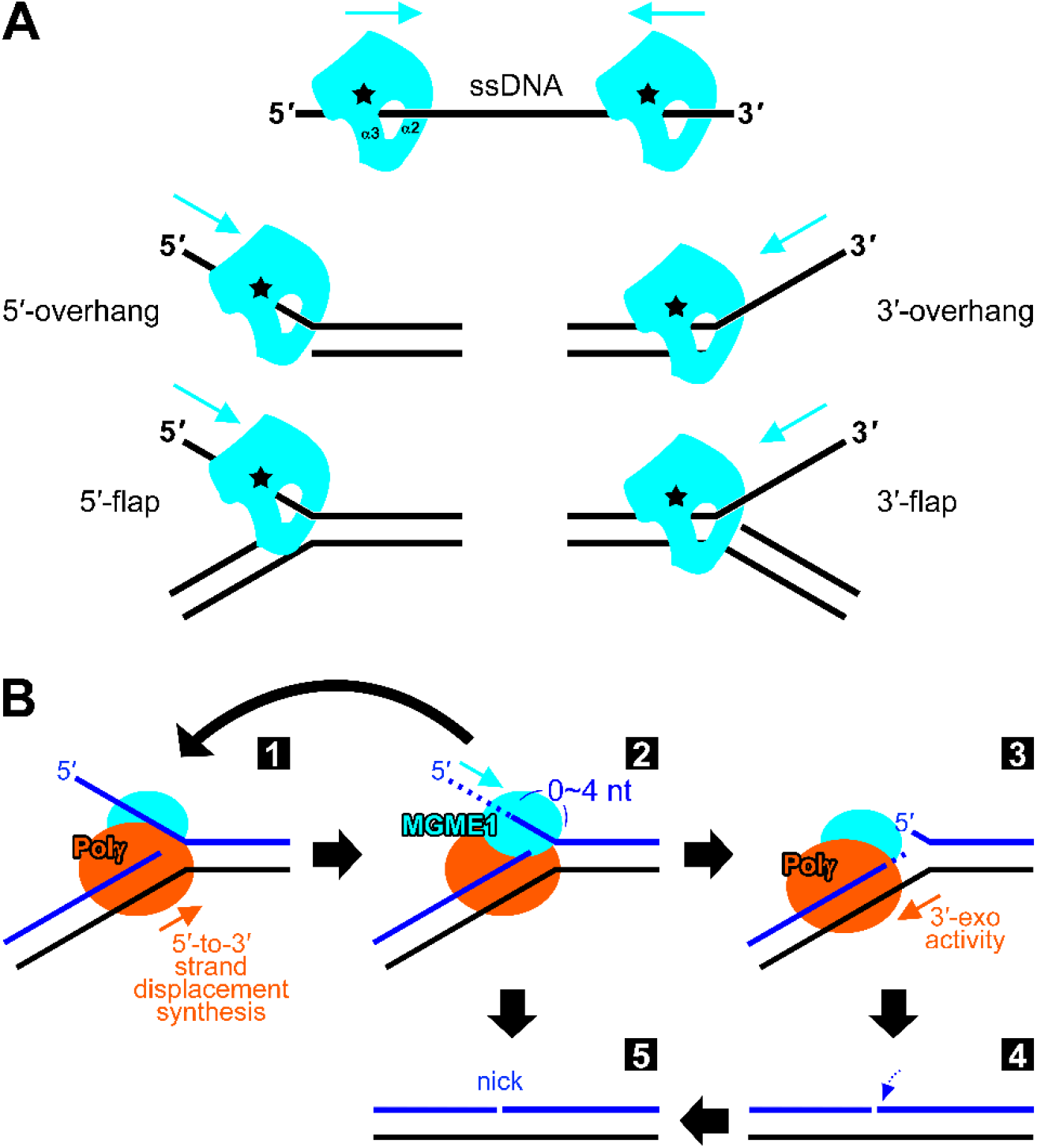
Working model of MGME1. (**A**) MGME1 exhibits bidirectional ss-specific exonuclease activity capable of removing ssDNA from various kinds of DNA duplexes. The catalytic site of MGME1 is labeled by a black star. (**B**) Cooperation between MGME1 and Polγ generates the ligatable nick at the finalization step of mtDNA replication. Upon encountering the 5′ end of the nascent DNA (blue), Polγ displaces the strand to generate a 5′ flap (step 1). Then, MGME1 trims the 5′ flap to fewer than 4 nucleotides depending on the GC content of the downstream duplex (step 2). Ligatable ends (step 5) may be readily generated at this point, or the process can repeat until the flap gets the opportunity to pair with the template strand (black) upon Polγ trimming the 3′ end of the nascent DNA (steps 3 and 4).

The essential 5′ processing of the nascent H-strand includes removal of the 2-nt RNA primer at CSB2, followed by digestion of ∼100 nucleotides of the DNA to OriH, and then production of a ligatable end at OriH to restore mtDNA integrity (1,2). This process does not seem to be solely attributable to a particular nuclease, since short-term depletion of any one of the known mitochondrial 5′ exonucleases, *i*.*e*., FEN1, DNA2 and EXOG, does not significantly perturb mtDNA copy number (38). However, cellular depletion of MGME1 alone, but not the other mitochondrial nucleases, led to 7S DNA accumulation, highlighting the specificity of MGME1 in 7S DNA metabolism (5-7,12,38). This scenario implies that 5′-end processing of 7S DNA may differ from that of the nascent H-strand. Nevertheless, how MGME1 participates in this particular process had been elusive. In terms of the molecular mechanism underlying nascent strand processing, our study provides insights into how MGME1 removes the 5′-flap to produce a ligatable nick to restore mtDNA integrity. Although the misalignment between the catalytic site and the ss-selective structural barrier of MGME1, *i*.*e*., the helical arch, prohibits the enzyme from cutting at the ss-ds junction, we found that MGME1 can unwind the downstream duplex in a GC content-dependent manner, explaining the imprecise cleavage around the ss-ds junction by MGME1 observed in the reported *in vitro* nuclease assays (5,7). Based on the model proposed by Uhler *et al*. (7) that illustrates collaboration between MGME1 and Polγ to generate a ligatable nick, we further detail this process by specifying the trimming range (within four nucleotides) of the two DNA ends, with one being shortened by MGME1 and the other being trimmed by the 3′-exo activity of Polγ (Figure 5). It should be noted that Polγ exhibits limited strand-displacement synthesis activity (6 to 9 nucleotides within 2 minutes, as reported previously (7,36)), as well as limited ability to trim the 3′ terminus (predominantly a single nucleotide in the absence of mismatches (39)). Accordingly, efficient generation of the ligatable nick (moving directly from step 2 to step 5 in Figure 5B) is determined by the sequence context of the duplex downstream of the ss-ds junction, with an AT-rich sequence facilitating the process.

Apart from their combinatory role in mtDNA synthesis, MGME1 and Polγ also work together to degrade mtDNA in response to DNA insults (21). In addition to evidence from mitochondrial-targeting restriction enzyme-triggered mtDNA degradation, depleting mtDNA from patient-derived fibroblasts lacking MGME1 by means of 2′,3′-dideoxycytidine (ddC) treatment resulted in delayed mtDNA breakdown, as well as impaired mtDNA recovery upon ddC withdrawal (5). This observation agrees with MGME1 exerting a dual role in maintaining mtDNA homeostasis. Furthermore, mice lacking MGME1 share common mtDNA aberrations with mice hosting Polγ deficient in 3′-exo activity (known as the mutator mice), including accumulations of linear subgenomic mtDNA fragments, mtDNA deletions and rearrangements (20,23,40). This scenario supports the tight functional coordination between MGME1 and Polγ’s 3′-exo activity. A model has been proposed to explain how these subgenomic fragments are generated in mice (41). However, it should be noted that the dual role of MGME1 and Polγ in mtDNA maintenance makes it complicated to dissect the mechanism giving rise to the mtDNA aberrations observed in genetically modified mouse models.

Deleting the dynamic N-terminus (residues 21-94) of MGME1 enhances both the enzyme’s 5′-exo and 3′-exo activity (Supplementary Figure S1F), suggesting that the N-terminus may exert an autoinhibitory role. In addition, it has been reported previously that overexpression of either WT (5,6) or catalytically-dead (D251N/K253A) (6) MGME1 in cultured cells induced 7S DNA depletion, implying that MGME1 may regulate 7S DNA levels through an unidentified function other than its nuclease activity. We anticipate that the dynamic N-terminus may contribute to the unidentified MGME1 function, including by regulating its nuclease activity and exerting a putative role in mediating protein-protein interactions as previously speculated by Yang *et. al*. (8). However, the entire N-terminus, which spans nearly one-third of the protein, has not been reported experimentally as forming an ordered or defined structure. The low per-residue confidence score (pLDDT) of the N-terminus in the AlphaFold2-predicted MGME1 model also reflects the dynamic nature of this region (Supplementary Figure S6A). Notably, there is a putative helix of about three turns (denoted αx herein) predicted for residues 48 to 60, which is located in a relatively conserved region of the N-terminus across MGME1 orthologs (Supplementary Figure S7). However, the position of αx relative to the enzyme core remains unclear due to the low pLDDT value of regions flanking that helix. Furthermore, we noticed that two regions that could not be resolved in our ΔN-MGME1-DNA structure, covering residues 107 to 118 and 190 to 202, are also mostly missing from other available MGME1 crystal structures (8), while they are predicted by AlphaFold2 to form ordered structures with moderate pLDDT scores (Supplementary Figure S6A). The surface representation of the AlphaFold2-predicted model reveals that these two regions form a continuous spine-like feature (Supplementary Figure S6B). Intriguingly, there is considerable residue variation in these regions among MGME1 orthologs (Supplementary Figure S7). Thus, how these features, *i*.*e*., the dynamic N-terminus and the non-conserved spine-like surface area, contribute to MGME1’s function remains to be fully explored.

By superimposing our resolved structure with the AlphaFold2-predicted model, we reconstituted a model of 5′-overhang DNA-bound MGME1 complex, with residues 95 to 130 of the enzyme forming intramolecular interactions with the protein surface (Supplementary Figure S6 B-E). Additional information concerning protein-DNA interactions can be gleaned from the model. For example, from the α3-side view, residue R127 can be identified as putatively interacting with the DNA backbone. This residue, along with R135 and the N-terminus of α1, forms a continuous and positively electrostatic groove to accommodate the negatively charged DNA backbone (Supplementary Figure S6D). Furthermore, the α2-side view reveals a positive electrostatic surface, which presumably facilitates DNA loading from this side of the enzyme. Note that some surface-exposed residues in our resolved structure could not be modeled due to a lack of respective electron density maps, such as K257 in the β3-α5 loop (Supplementary Figure S6E). Thus, the AlphaFold2-predicted model serves as a good complement to our experimentally determined structure, enabling more comprehensive structural analyses.

Taken together, in this report, we provide structural insights into the DNA 5′-end processing activity of MGME1, which serves as an ancillary 5′ exonuclease alongside Polγ in maintaining mtDNA integrity. MGME1 operates as a rigid clamp that slides along DNA by means of a ss-selective helical arch structure located downstream of its catalytic site. Despite the gap between the catalytic site and this arch, MGME1 can partially unwind the duplex downstream of the ss-ds junction, helping it to better resolve a 5′-ssDNA flap. Given the dynamic nature of MGME1’s N-terminus, which contrasts with its rigid core, we hypothesize that the N-terminus exerts a regulatory role. Whether this particular region contributes to MGME1’s interplay with Polγ to maintain mtDNA warrants further investigation.

## DATA AVAILABILITY

The coordinate and structure factor of the resolved crystal structure of the ΔN-MGME1-5′-overhang DNA complex has been deposited to the Protein Data Bank with accession code 8XA9.

## SUPPLEMENTARY DATA

Supplementary Data are available at bioRxiv Online.

## ACKNOWLEDGMENTS

Portions of this research were carried out at the National Synchrotron Radiation Research Center, a national user facility supported by the National Science and Technology Council of Taiwan. The Synchrotron Radiation Protein Crystallography Facility is supported by the National Core Facility Program for Biotechnology. We thank Dr. Hanna S. Yuan and Dr. Nei-Li Chan for their suggestions during preparation of the manuscript.

## AUTHOR CONTRIBUTIONS

E.Y.C.M. designed and performed the experiments, including protein purification, as well as crystallization, biochemical and mutagenesis analyses. H.Y.Y. contributed to the initial crystallization screening using full-length MGME1. C.C.W. and E.Y.C.M. collected the X-ray diffraction data and determined the structure. E.Y.C.M. analyzed the data and generated the figures. C.C.W. wrote the manuscript.

## REFERENCES

1. Uhler, J.P. and Falkenberg, M. (2015) Primer removal during mammalian mitochondrial DNA replication. DNA Repair, 34, 28–38.

2. Gustafsson, C.M., Falkenberg, M. and Larsson, N.G. (2016) Maintenance and Expression of Mammalian Mitochondrial DNA. Annu. Rev. Biochem., 85, 133–160.

3. Suomalainen, A. and Battersby, B.J. (2018) Mitochondrial diseases: the contribution of organelle stress responses to pathology. Nat. Rev. Mol. Cell Biol., 19, 77–92.

4. Rahman, S. and Copeland, W.C. (2019) POLG-related disorders and their neurological manifestations. Nat. Rev. Neurol., 15, 40–52.

5. Kornblum, C., Nicholls, T.J., Haack, T.B., Schöler, S., Peeva, V., Danhauser, K., Hallmann, K., Zsurka, G., Rorbach, J., Iuso, A. et al. (2013) Loss-of-function mutations in MGME1 impair mtDNA replication and cause multisystemic mitochondrial disease. Nat. Genet., 45, 214–219.

6. Szczesny, R.J., Hejnowicz, M.S., Steczkiewicz, K., Muszewska, A., Borowski, L.S., Ginalski, K. and Dziembowski, A. (2013) Identification of a novel human mitochondrial endo-/exonuclease Ddk1/c20orf72 necessary for maintenance of proper 7S DNA levels. Nucleic Acids Res., 41, 3144–3161.

7. Uhler, J.P., Thörn, C., Nicholls, T.J., Matic, S., Milenkovic, D., Gustafsson, C.M. and Falkenberg, M. (2016) MGME1 processes flaps into ligatable nicks in concert with DNA polymerase γ during mtDNA replication. Nucleic Acids Res., 44, 5861–5871.

8. Yang, C., Wu, R., Liu, H., Chen, Y., Gao, Y., Chen, X., Li, Y., Ma, J., Li, J. and Gan, J. (2018) Structural insights into DNA degradation by human mitochondrial nuclease MGME1. Nucleic Acids Res., 46, 11075–11088.

9. Misic, J., Milenkovic, D., Al-Behadili, A., Xie, X., Jiang, M., Jiang, S., Filograna, R., Koolmeister, C., Siira, S.J., Jenninger, L. et al. (2022) Mammalian RNase H1 directs RNA primer formation for mtDNA replication initiation and is also necessary for mtDNA replication completion. Nucleic Acids Res., 50, 8749–8766.

10. Rocha, E.B.D., Rodrigues, K.D., Montouro, L.A.M., Coelho, E.N., Kouyoumdjian, J.A., Kok, F., N ó brega, P.R., Graca, C.R., Morita, M.D.A. and Estephan, E.D. (2023) A case of mitochondrial DNA depletion syndrome type 11-expanding the genotype and phenotype. Neuromuscular Disord., 33, 692–696.

11. Nicholls, T.J. and Minczuk, M. (2014) In D-loop: 40 years of mitochondrial 7S DNA. Exp. Gerontol., 56, 175–181.

12. Nicholls, T.J., Zsurka, G., Peeva, V., Schöler, S., Szczesny, R.J., Cysewski, D., Reyes, A., Kornblum, C., Sciacco, M., Moggio, M. et al. (2014) Linear mtDNA fragments and unusual mtDNA rearrangements associated with pathological deficiency of MGME1 exonuclease. Hum. Mol. Genet., 23, 6147–6162.

13. Lima, W.F., Rose, J.B., Nichols, J.G., Wu, H.J., Migawa, M.T., Wyrzykiewicz, T.K., Siwkowski, A.M. and Crooke, S.T. (2007) Human RNase H1 discriminates between subtle variations in the structure of the heteroduplex substrate. Mol. Pharmacol., 71, 83–91.

14. Nowotny, M., Gaidamakov, S.A., Ghirlando, R., Cerritelli, S.M., Crouch, R.J. and Yang, W. (2007) Structure of human RNase h1 complexed with an RNA/DNA hybrid: Insight into HIV reverse transcription. Mol. Cell, 28, 264–276.

15. Cerritelli, S.M. and Crouch, R.J. (2009) Ribonuclease H: the enzymes in eukaryotes. Febs. J., 276, 1494–1505.

16. Kalifa, L., Beutner, G., Phadnis, N., Sheu, S.S. and Sia, E.A. (2009) Evidence for a role of FEN1 in maintaining mitochondrial DNA integrity. DNA Repair, 8, 1242–1249.

17. Al-Behadili, A., Uhler, J.P., Berglund, A.K., Peter, B., Doimo, M., Reyes, A., Wanrooij, S., Zeviani, M. and Falkenberg, M. (2018) A two-nuclease pathway involving RNase H1 is required for primer removal at human mitochondrial OriL. Nucleic Acids Res., 46, 9471–9483.

18. Zheng, L., Zhou, M.A., Guo, Z.G., Lu, H.M., Qian, L.M., Dai, H.F., Qiu, J.Z., Yakubovskaya, E., Bogenhagen, D.F., Demple, B. et al. (2008) Human DNA2 Is a Mitochondrial Nuclease/Helicase for Efficient Processing of DNA Replication and Repair Intermediates. Mol. Cell, 32, 325–336.

19. Wu, C.C., Lin, J.L., Yang-Yen, H.F. and Yuan, H.S. (2019) A unique exonuclease ExoG cleaves between RNA and DNA in mitochondrial DNA replication. Nucleic Acids Res., 47, 5405–5413.

20. Nissanka, N., Bacman, S.R., Plastini, M.J. and Moraes, C.T. (2018) The mitochondrial DNA polymerase gamma degrades linear DNA fragments precluding the formation of deletions. Nat. Commun., 9.

21. Peeva, V., Blei, D., Trombly, G., Corsi, S., Szukszto, M.J., Rebelo-Guiomar, P., Gammage, P.A., Kudin, A.P., Becker, C., Altmuller, J. et al. (2018) Linear mitochondrial DNA is rapidly degraded by components of the replication machinery. Nat. Commun., 9, 1727.

22. Medeiros, T.C. and Graef, M. (2019) Autophagy determines mtDNA copy number dynamics during starvation. Autophagy, 15, 178–179.

23. Matic, S., Jiang, M., Nicholls, T.J., Uhler, J.P., Dirksen-Schwanenland, C., Polosa, P.L., Simard, M.L., Li, X.P., Atanassov, I., Rackham, O. et al. (2018) Mice lacking the mitochondrial exonuclease MGME1 accumulate mtDNA deletions without developing progeria. Nat. Commun., 9.

24. GraphPad Software, B., Massachusetts USA, http://www.graphpad.com.

25. Otwinowski, Z. and Minor, W. (1997) Processing of X-ray diffraction data collected in oscillation mode. Method Enzymol., 276, 307–326.

26. Jumper, J., Evans, R., Pritzel, A., Green, T., Figurnov, M., Ronneberger, O., Tunyasuvunakool, K., Bates, R., Zídek, A., Potapenko, A. et al. (2021) Highly accurate protein structure prediction with AlphaFold. Nature, 596, 583–589.

27. Varadi, M., Anyango, S., Deshpande, M., Nair, S., Natassia, C., Yordanova, G., Yuan, D., Stroe, O., Wood, G., Laydon, A. et al. (2022) AlphaFold Protein Structure Database: massively expanding the structural coverage of protein-sequence space with high-accuracy models. Nucleic Acids Res., 50, D439–D444.

28. Bunkóczi, G., Echols, N., McCoy, A.J., Oeffner, R.D., Adams, P.D. and Read, R.J. (2013) Phaser.MRage: automated molecular replacement. Acta Crystallogr. D., 69, 2276–2286.

29. Adams, P.D., Afonine, P.V., Bunkóczi, G., Chen, V.B., Davis, I.W., Echols, N., Headd, J.J., Hung, L.W., Kapral, G.J., Grosse-Kunstleve, R.W. et al. (2010) PHENIX: a comprehensive Python-based system for macromolecular structure solution. Acta Crystallogr. D., 66, 213–221.

30. Emsley, P., Lohkamp, B., Scott, W.G. and Cowtan, K. (2010) Features and development of Coot. Acta Crystallogr. D., 66, 486–501.

31. Afonine, P.V., Grosse-Kunstleve, R.W., Echols, N., Headd, J.J., Moriarty, N.W., Mustyakimov, M., Terwilliger, T.C., Urzhumtsev, A., Zwart, P.H. and Adams, P.D. (2012) Towards automated crystallographic structure refinement with phenix.refine. Acta Crystallogr. D., 68, 352–367.

32. The PyMOL Molecular Graphics System, V.S., LLC.

33. Urrutia, K.M., Xu, W.Y. and Zhao, L.L. (2022) The 5′-phosphate enhances the DNA-binding and exonuclease activities of human mitochondrial genome maintenance exonuclease 1 (MGME1). J. Biol. Chem., 298.

34. Tsutakawa, S.E., Classen, S., Chapados, B.R., Arvai, A.S., Finger, L.D., Guenther, G., Tomlinson, C.G., Thompson, P., Sarker, A.H., Shen, B.H. et al. (2011) Human Flap Endonuclease Structures, DNA Double-Base Flipping, and a Unified Understanding of the FEN1 Superfamily. Cell, 145, 198–211.

35. Tsutakawa, S.E., Thompson, M.J., Arvai, A.S., Neil, A.J., Shaw, S.J., Algasaier, S.I., Kim, J.C., Finger, L.D., Jardine, E., Gotham, V.J.B. et al. (2017) Phosphate steering by Flap Endonuclease 1 promotes 5′-flap specificity and incision to prevent genome instability. Nat Commun., 8.

36. Macao, B., Uhler, J.P., Siibak, T., Zhu, X.F., Shi, Y.H., Sheng, W.W., Olsson, M., Stewart, J.B., Gustafsson, C.M. and Falkenberg, M. (2015) The exonuclease activity of DNA polymerase γ is required for ligation during mitochondrial DNA replication. Nat. Commun., 6.

37. Torregrosa-Muñumer, R., Hangas, A., Goffart, S., Blei, D., Zsurka, G., Griffith, J., Kunz, W.S. and Pohjoismäki, J.L.O. (2019) Replication fork rescue in mammalian mitochondria. Sci. Rep., 9.

38. Moretton, A., Morel, F., Macao, B., Lachaume, P., Ishak, L., Lefebvre, M., Garreau-Balandier, I., Vernet, P., Falkenberg, M. and Farge, G. (2017) Selective mitochondrial DNA degradation following double-strand breaks. Plos One, 12.

39. He, Q., Shumate, C.K., White, M.A., Molineux, I.J. and Yin, Y.W. (2013) Exonuclease of human DNA polymerase gamma disengages its strand displacement function. Mitochondrion, 13, 592–601.

40. Milenkovic, D., Sanz-Moreno, A., Calzada-Wack, J., Rathkolb, B., Amarie, O.V., Gerlini, R., Aguilar-Pimentel, A., Misic, J., Simard, M.L.S., Wolf, E.T. et al. (2022) Mice lacking the mitochondrial exonuclease MGME1 develop inflammatory kidney disease with glomerular dysfunction. Plos Genet., 18.

41. Falkenberg, M. and Gustafsson, C.M. (2020) Mammalian mitochondrial DNA replication and mechanisms of deletion formation. Crit. Rev. Biochem. Mol., 55, 509–524.

