## Supplementary material for "Structural Basis of How MGME1 Processes DNA 5′ Ends to Maintain Mitochondrial Genome Integrity": MGME1 MS_NARv_SI_1207.pdf

**Supplementary Information**
**Of**

### Structural Basis of How MGME1 Processes DNA 5′ Ends to Maintain Mitochondrial Genome Integrity

**Authors:**

Eric Y.C. Mao, Han-Yi Yen, and Chyuan-Chuan Wu[*]

[*]Corresponding author

Corresponding

This file contains:

Supplementary Table S1-S2

Supplementary Figure S1-S7

Supplementary References 1-6

### SUPPLEMENTARY TABLES

**Supplementary Table S1.** Data collection and refinement parameters of X-ray crystallography

| Structure name | 5′-overhang DNA-bound MGME1 |
|---|---|
| PDB ID | 8XA9 |
| Data collection | |
| Resolution range (Å) | 30.00 – 2.35<br>(2.40 – 2.35) |
| Space group | C 1 2 1 |
| Unit cell (Å, degree) | a=180.756, b=56.193, c=114.445,<br>$\alpha$=90°, $\beta$=113.023°, $\gamma$=90° |
| Total reflections | 143,955 |
| Unique reflections | 43,737 (2,393) |
| Multiplicity | 3.3 (3.1) |
| Completeness (%) | 96.4 (95.5) |
| Mean I/sigma(I) | 19.17 (2.39) |
| Wilson B-factor | 48.47 |
| R-merge<br>R-meas<br>R-pim<br>CC1/2 | 0.056 (0.504)<br>0.066 (0.608)<br>0.035 (0.335)<br>(0.925) |
| Refinement | |
| Reflections used in refinement<br>Reflections used for R-free | 43,703 (3,563)<br>2,162 (170) |
| R-work | 0.2579 (0.3420) |
| R-free | 0.2778 (0.3423) |
| RMS (bonds, Å) | 0.004 |
| RMS (angles, degree) | 0.62 |
| Ramachandran favoured (%)<br>Ramachandran allowed (%)<br>Ramachandran outliers (%) | 96.39<br>3.61<br>0.00 |
| Rotamer outliers (%)<br>Average B-factor<br>Clash score | 0.00<br>60.89<br>3.09 |

Statistics for the highest-resolution shell are shown in parentheses.

**Supplementary Table S2.** Sequence of the synthetic oligonucleotides used in this study

| Name | Length | Sequence (5′→3′) | Figure |
|---|---|---|---|
| **The 7-nt 5′-overhang DNA for crystallization** | | | |
| Scissile strand | 18 | TTTTTTTGCTGGCGGTCG | 1 |
| Non-scissile strand | 11 | CGACCGCCAGC | 1 |
| **Fluorescence-labeled probes** | | | |
| 5'P-58nt-HighGC-3'Cy5 | 58 | 5'P-TTTTTTTTTTTTTTTTTTTTTTTTTTTTTTTTTTTTTTT TTGGCGACGGCAGCGAGGC-**3'Cy5** | 2A, 3A |
| 5'FAM-58nt | 58 | **5'FAM-**TGGCGACGGCAGCGAGGC TTTTTTTTTTTTTTTTTTTTTTTTTTTTTTTTTTTTTTTT | 2B |
| 5'P-58nt-(iCy3)$_2$-3'Cy5 | 58 | 5'P-TTTTTTTTTT**T(Cy3)**TTTTTTTTTTTTTTTTTTT **T(Cy3)**TTTTTTTTTTGGCGACGGCAGCGAGGC**-3'Cy5** | 2E |
| 5'Cy5-58nt-(iCy3)$_2$ | 58 | **5'Cy5**-TGGCGACGGCAGCGAGGCTTTTTTTTT **T(Cy3)**TTTTTTTTTTTTTTTTTTT**T(Cy3)**TTTTTTTTTT | 2E |
| 5'P-58nt-LowGC-3'Cy5 | 58 | 5'P-TTTTTTTTTTTTTTTTTTTTTTTTTTTTTTTTTTTTTTT TTAATAACGGCAGCGAGGC-**3'FAM** | 3B |
| 5'P-25nt-HighGC-3FAM | 25 | 5'P-GTCTAACGCTGACTCGCTACGTACC-**3'FAM** | 3C, 3E, 4A, 4C, 4E, 4G, 4H |
| 5P-25nt-LowGC-3FAM | 25 | 5'P-GTCTAACTAATAACGGCAGCGAGGC-**3'FAM** | 3D, 3E |
| 5'FAM-25nt | 25 | **5'FAM**-CCATGCATCGCTCAGTCGCAATCTG | 4B, 4D |
| 5'FAM-22nt | 22 | 5'P-TAACGCTGACTCGCTACGTACC-**3'FAM** | 4F, 4I |
| **Fluorescence-labeled markers (M)** | | | |
| HighGC 58nt-3'Cy5- M1 | 22 | 5'P-TTTTTGGCGACGGCAGCGAGGC-**3'Cy5** | 2A, 3A |
| HighGC 58nt-3'Cy5-M2 | 18 | 5'P-TGGCGACGGCAGCGAGGC-**3'Cy5** | 2A, 2E, 3A |
| HighGC 5'FAM-58nt-M1 | 18 | **5'FAM**-TGGCGACGGCAGCGAGGC | 2B |
| HighGC 5'FAM-58nt-M2 | 15 | **5'FAM**-TGGCGACGGCAGCGA | 2B |
| LowGC-58nt-Cy5-M | 18 | 5'P-TAATAACGGCAGCGAGGC-**3'Cy5** | 3B, 3D |
| HighGC-25nt-FAM-M | 18 | 5'P-GCTGACTCGCTACGTACC-**3'FAM** | 3C |
| **Label-free complementary (C) oligonucleotides** | | | |
| C-strand for 40nt overhang | 18 | GCCTCGCTGCCGTCGCCA | 2A and 2B |
| 38nt C-strand for HighGC-40nt flap | 38 | GCCTCGCTGCCGTCGCCACGCTTTGTAGGCAGGTACGC | 3A |
| 38nt C-strand for LowGC-40 and 7nt flap | 38 | GCCTCGCTGCCGTTATTACGCTTTGTAGGCAGGTACGC | 3B, 3D, 3E |
| 38nt C-strand for HighGC-7nt flap | 38 | GGTACGTAGCGAGTCAGCCGCTTTGTAGGCAGGTACGC | 3C, 3E |
| 20nt C-strand for flap | 20 | GCGTACCTGCCTACAAAGCG | 3A-3E |
| C-strand for 7nt 5'-overhang DNA | 18 | GGTACGTAGCGAGTCAGC | 4A, 4C, 4E-I |
| C-strand for 7nt 3'-overhang DNA | 18 | CGACTGAGCGATGCATGG | 4B, 4D |
| **Primers for site-directed mutagenesis#** | | | |
| F147A_Forward | CAACAGGTT**GC**CTTGTTGGAGAGGTGG | | N/A |
| F147A_Reverse | CCAACAAG**GC**AACCTGTTGTTTTGTCATGG | | |
| W152A_ Forward | TTGGAGAGG**GC**GAAACAGCGGATGATTCTGGAAC | | |
| W152A_ Reverse | CGCTGTTTC**GC**CCTCTCCAACAAGAAAACCTG | | |
| F173A_ Forward | CAAACGTC**GC**TTTACAAGGGAAACGG | | |
| F173A_ Reverse | CCCTTGTAAA**GC**GACGTTTGAAGTGTATTC | | |
| F261A_ Forward | CCAAAGCCT**GC**TATTCAAAGTACATTTGACAACCC | | |
| F261A_ Reverse | CTTTGAATA**GC**AGGCTTTGGTTTCTCTGATGTC | | |
| F266A_ Forward | CAAAGTACA**GC**TGACAACCCACTGCAAG | | |
| F266A_ Reverse | GGTTGTCA**GC**TGTACTTTGAATAAAAGGC | | |

#The targeted cordon is underlined with the mutated nucleotides shown in bold

### SUPPLEMENTARY FIGURES

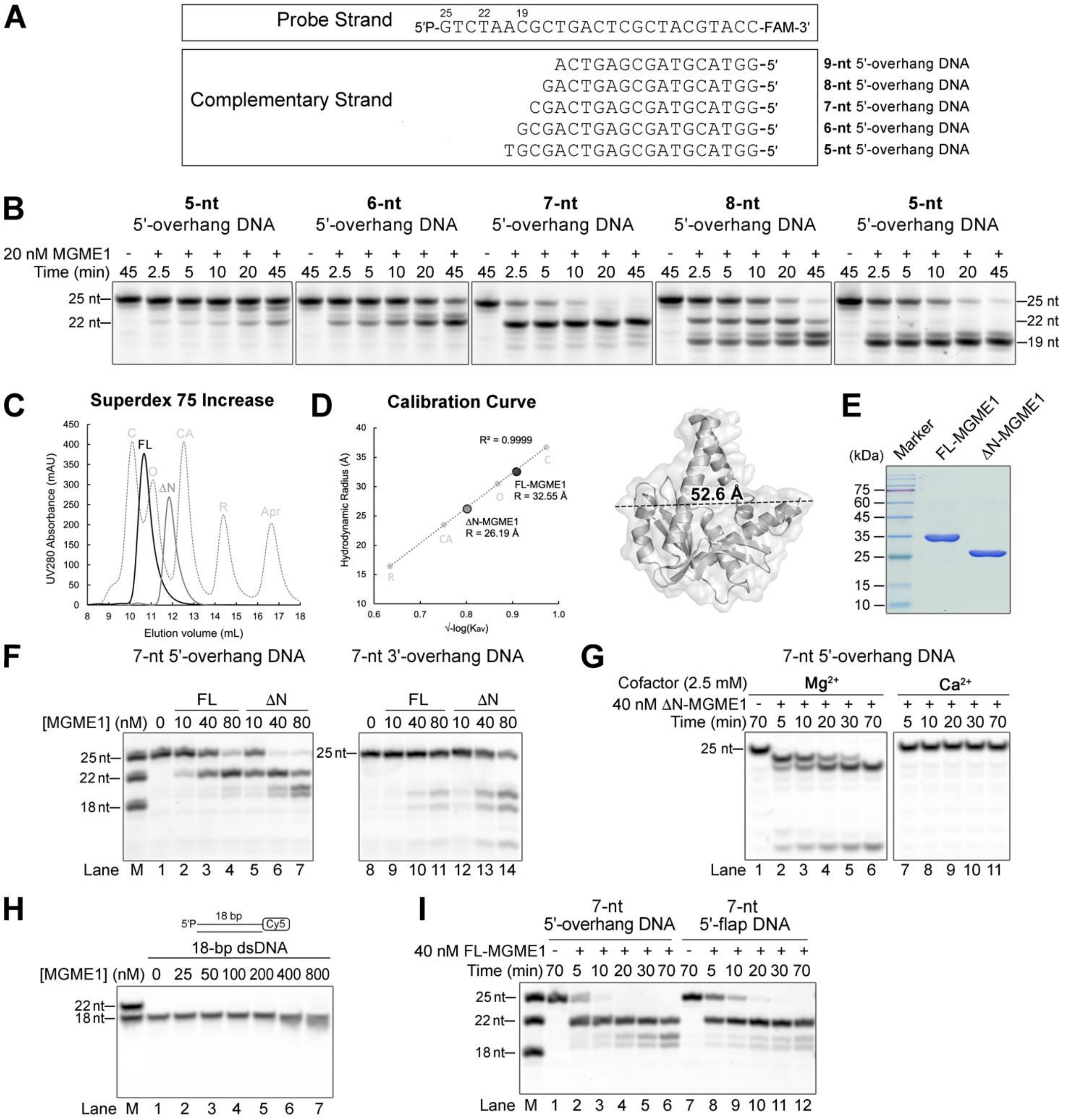


**Supplementary Figure S1.** Purification and nuclease activity test of full-length (FL-) and ΔN-MGME1. (**A**) The 5′-overhang DNA duplexes used for the assay. The substrates were prepared by annealing the 25-nt probe strand with different complementary strands to generate 5′-ssDNA overhangs ranging from 5 to 9 nt as shown in **B**. (**B**) FL-MGME1 (residues 21-344) in digesting 5′-overhang DNA duplexes. (**C**) Size-exclusion chromatogram (SEC) of the purified FL- and ΔN-MGME1. The dotted curve shows the profile of standard proteins (Cytiva): conalbumin (C; 36.7 Å) (1), ovalbumin (O; 30.5 Å), carbonic anhydrase (CA; 23.5 Å) (2), and ribonuclease A (R; 16.4 Å). (**D**) The calibration curve of the SEC analysis as shown in **C**. (**E**) SDS-PAGE of the purified FL and ΔN-MGME1. The theoretical molecular weights of FL- and ΔN-MGME1 are 37.0 and 29.0 kDa, respectively. (**F**) ΔN-MGME1 exhibits a higher nuclease activity compared to FL-MGME1. The reactions were incubated at 37°C for 10 min. (**G**) $Ca^{2+}$ serves as an inhibitory ion for MGME1. (**H**) MGME1 could not digest dsDNA unless at a high concentration. 100 nM dsDNA substrate was incubated with the enzyme at 37°C for 10 min. Notable degradation was only observed at high MGME1 concentration (>4 fold of substrate concentration). (**I**) FL-MGME1 generated the same DNA cleavage pattern from 5′-overhang or 5′-flap DNA

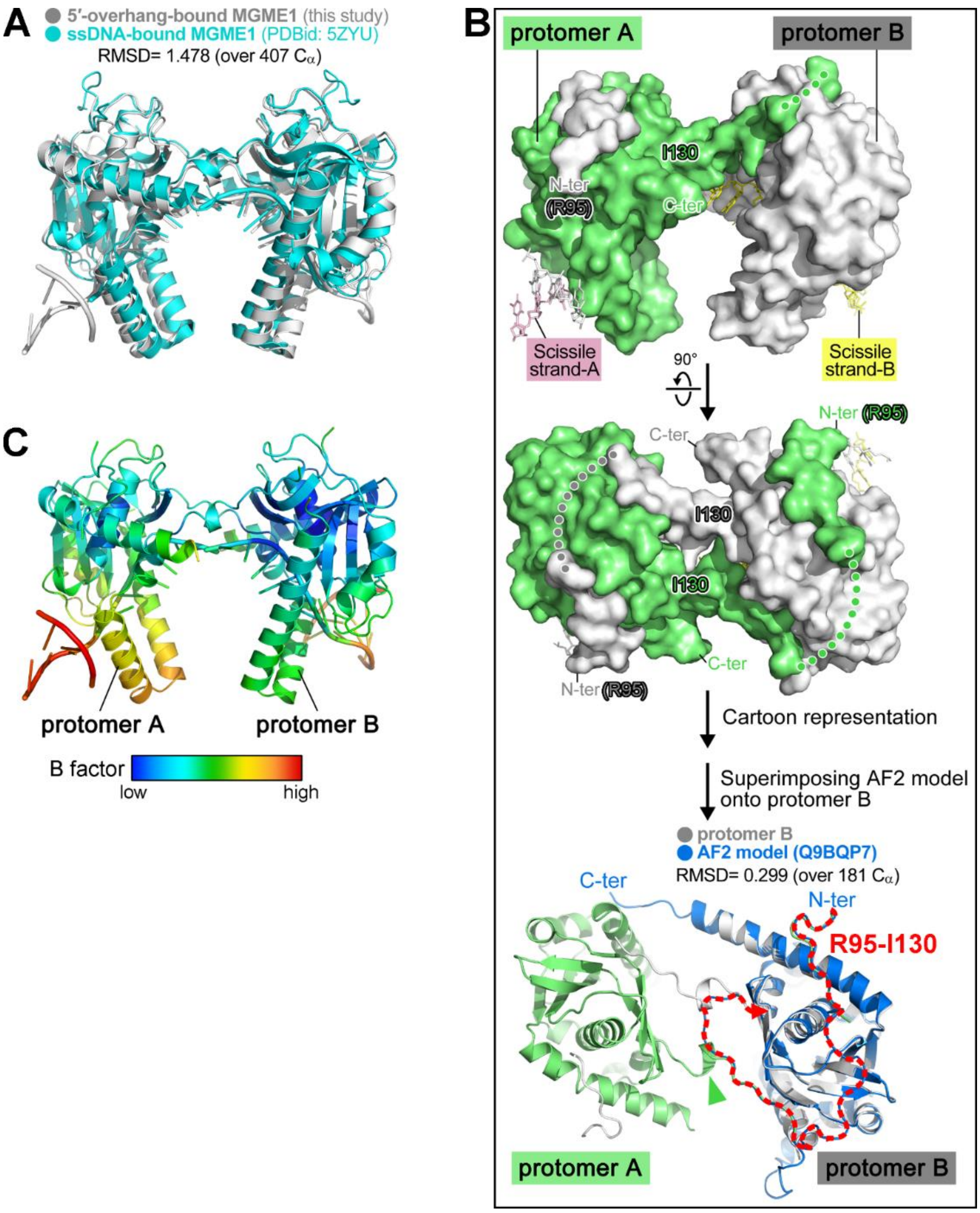


**Supplementary Figure S2. The swapped N-terminus in the crystal structures of MGME1.** (**A**) Superimposition of the 5′-overhang DNA-bound (this study) and ssDNA-bound (PDB ID: 5ZYU) MGME1 structures. All residues in an asymmetry unit were aligned by secondary structure in PyMOL (3), yielding a root-mean-square-deviation (RMSD) value of 1.478 (over 407 Cα). Alignment using one of the protomers yielded an RMSD of 0.359 (over 192 Cα) as shown in Figure 1C. (**B**) Surface representation of the two protomers in the 5′-overhang DNA-bound MGME1 structure. Missing regions are illustrated by dotted lines. In the lowest panel, the AlphaFold2 (AF2)-predicted MGME1 model (UniProt session number: Q9BQP7, shown in blue) (4,5) was aligned to the protomer B (RMSD=0.299 (over 181 Cα)). The red dashed arrow illustrates the main-chain track of residues 95 to 130 (R95-I130) in the AF2 model. The green arrowhead indicates the turning point of the N-terminus of protomer A, which protrudes toward and packs against protomer B in the same way as residues 95 to 130 of the AF2 model. (**C**) The two protomers of the 5′-overhang DNA-bound MGME1 structure colored by B factor.

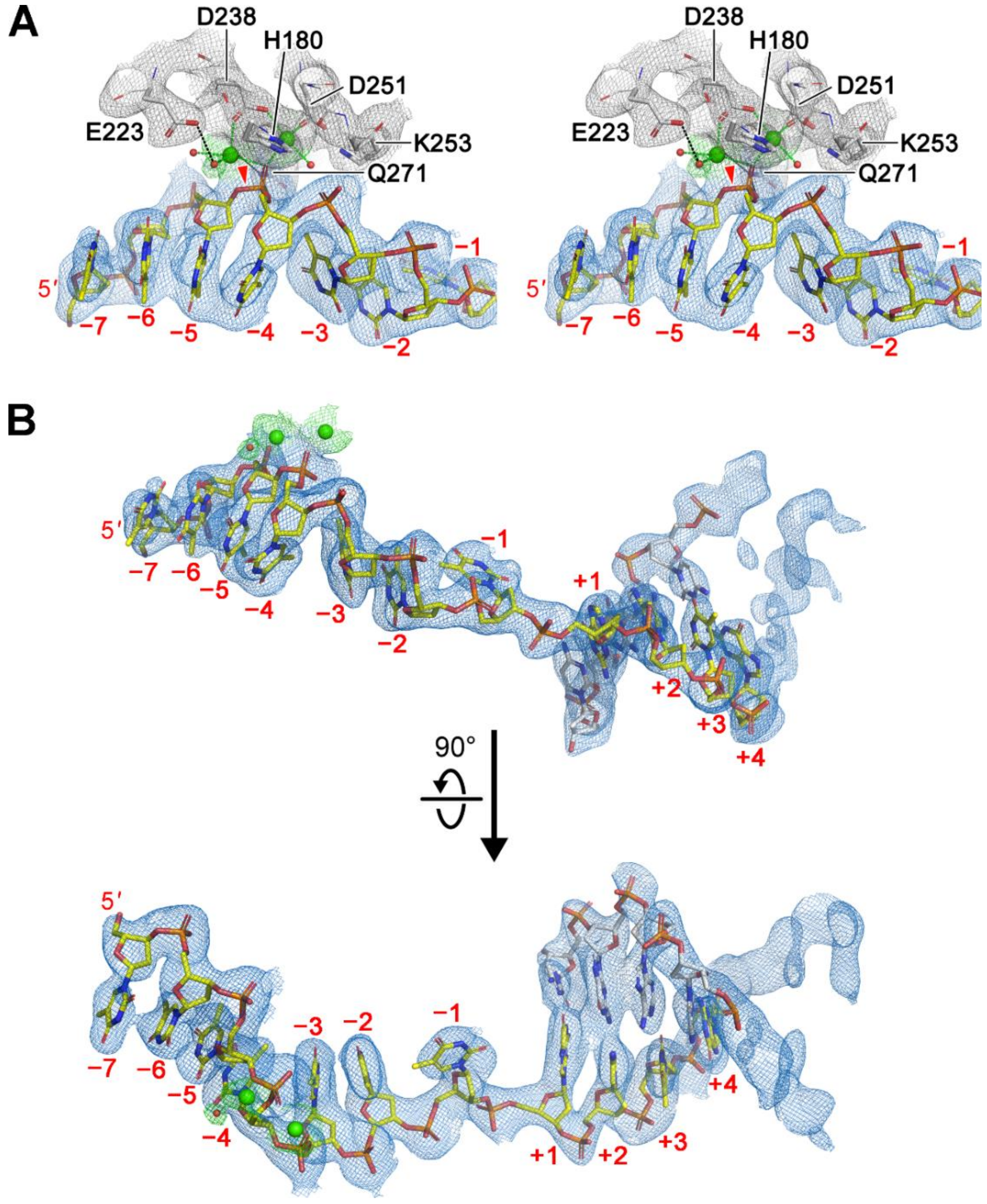


**Supplementary Figure S3.** The composition omitted maps of the bound 5′-overhang DNA in the complex formed by chains B, C and D in our solved structure. (**A**) Stereo view (wall eye) of the $Ca^{2+}$-stabilized pre-cleavage state of the catalytic site. (**B**) The DNA maps. For all panels, the catalytic-site-chelated $Ca^{2+}$ ions are shown as green spheres. Waters are shown as red spheres. $Ca^{2+}$ coordinations are illustrated as green dotted lines. Black dotted lines show the water-mediated interactions between $Ca^{2+}$ and protein residues.

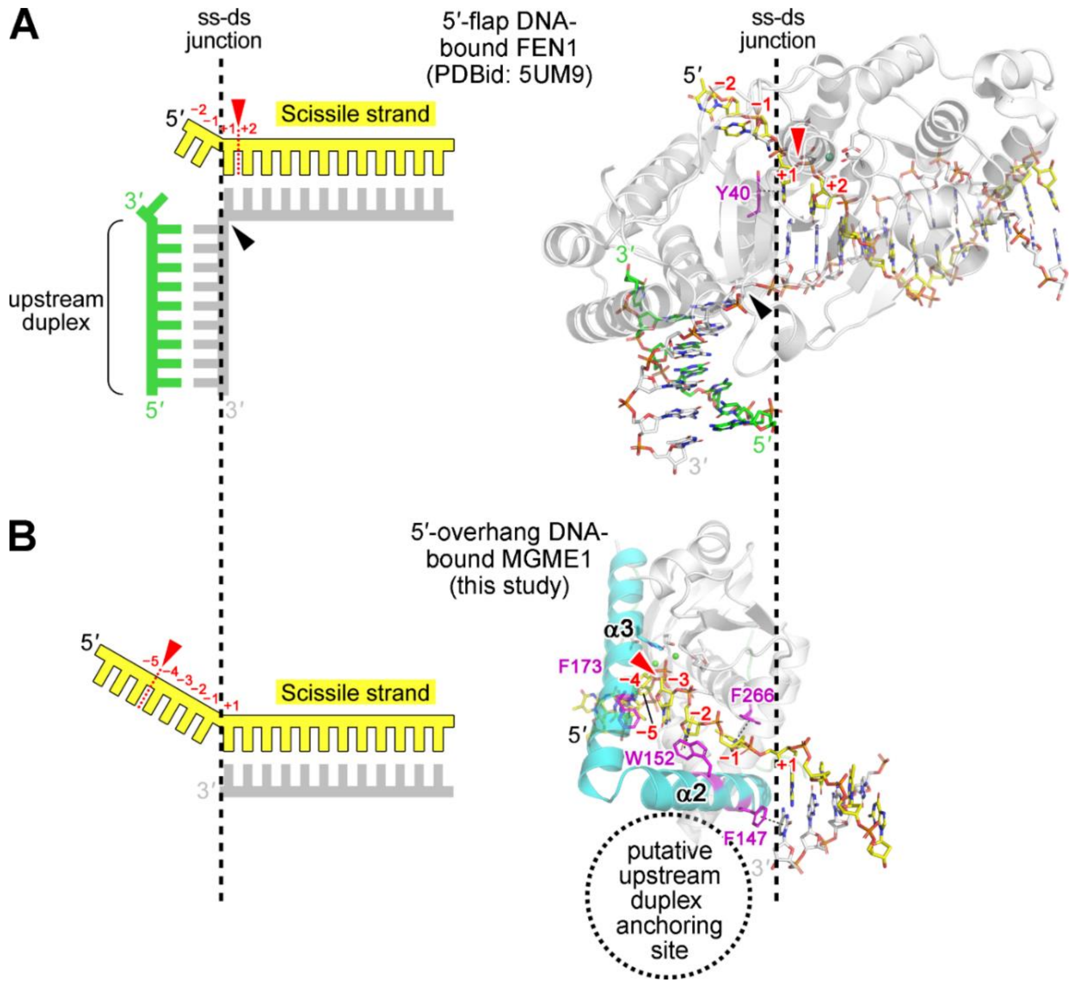


**Supplementary Figure S4.** Comparing the DNA-bound structures of MGME1 and FEN1. (**A**) The 5′-flap DNA-bound FEN1 structure (PDB ID: 5UM9) (6). Protein is shown in the transparent cartoon representation and DNA in the stick representation. The non-scissile strand is shown in grey and the paired upstream strand is in green. The black arrowheads indicate the kink of the non-scissile strand. The samarium ($Sm^{3+}$) bound in the catalytic site is shown as a sphere. The aromatic residue, Y40, stacks with the +1 nucleotide is shown and highlighted in magenta. (**B**) The 5′-overhang DNA-bound MGME1 structure (this study). The structure is shown and colored according to Figure 1. Cartoon representations of the bound substrates are shown on the left for both structures. The black dotted lines for all panels illustrate the stacking between the bound nucleotide and the selected aromatic residues. Red arrowheads indicate the cleavage sites of the enzymes.

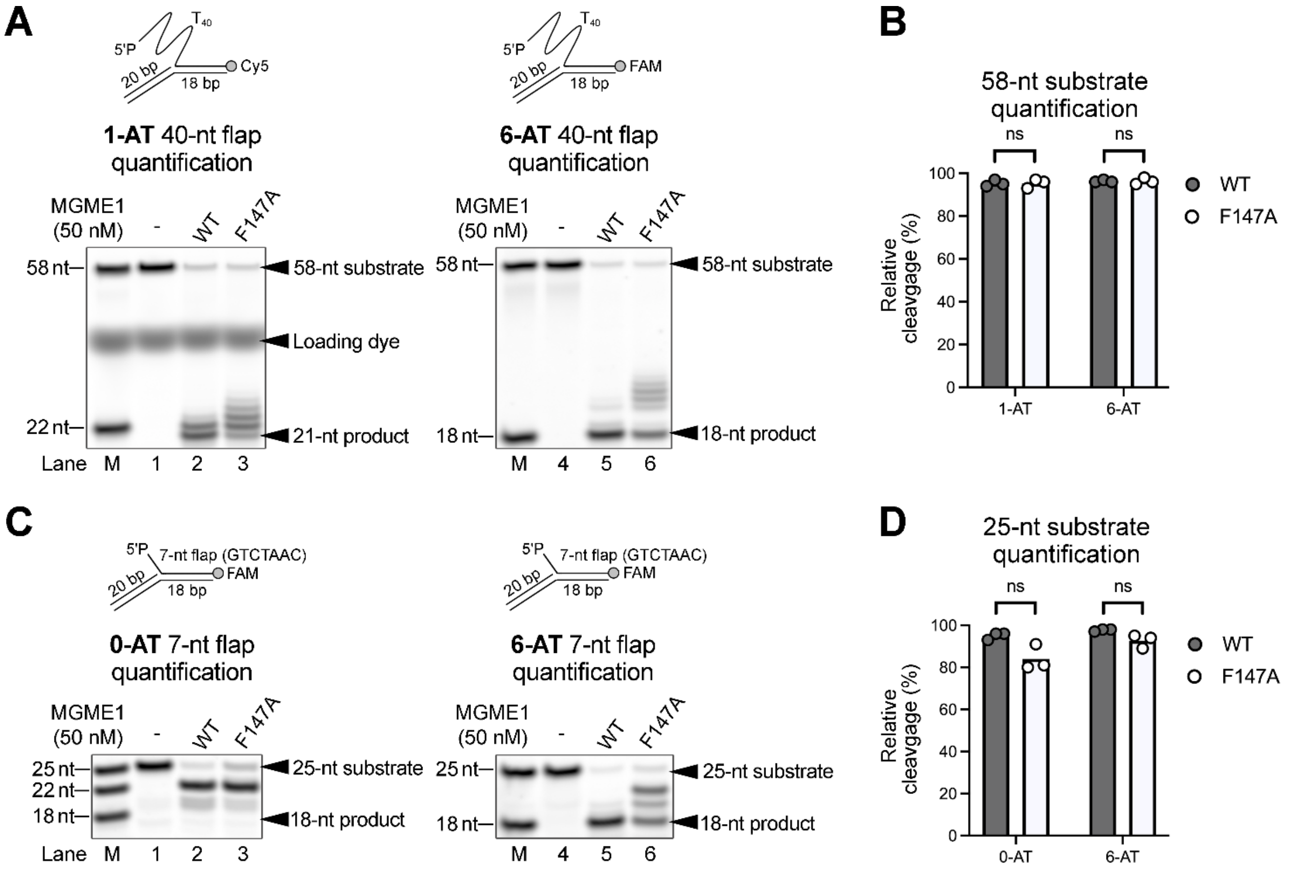


**Supplementary Figure S5.** Quantification of MGME1-derived DNA cleavage products as shown in Figure 3. (**A**) Activity assay of MGME1 in digesting 40-nt 5′-flap (1-AT and 6-AT) with an incubation time of 20 min. (**B**) Quantification of the 58-nt substrates in (**A**) shows no significant difference in the activity between WT-MGME1 and MGME1-F147A in digesting 40-nt 5′-flap. (**C**) Activity assay of MGME1 in digesting 7-nt 5′-flap (0-AT and 6-AT) with an incubation time of 20 min. (**D**) Quantification of the 25-nt substrates in (**C**) shows no significant difference in activity between WT-MGME1 and MGME1-F147A in digesting 7-nt 5′-flap. Two-tailed paired t-test was used to compare the mean values from two populations, ns: statistically non-significant.

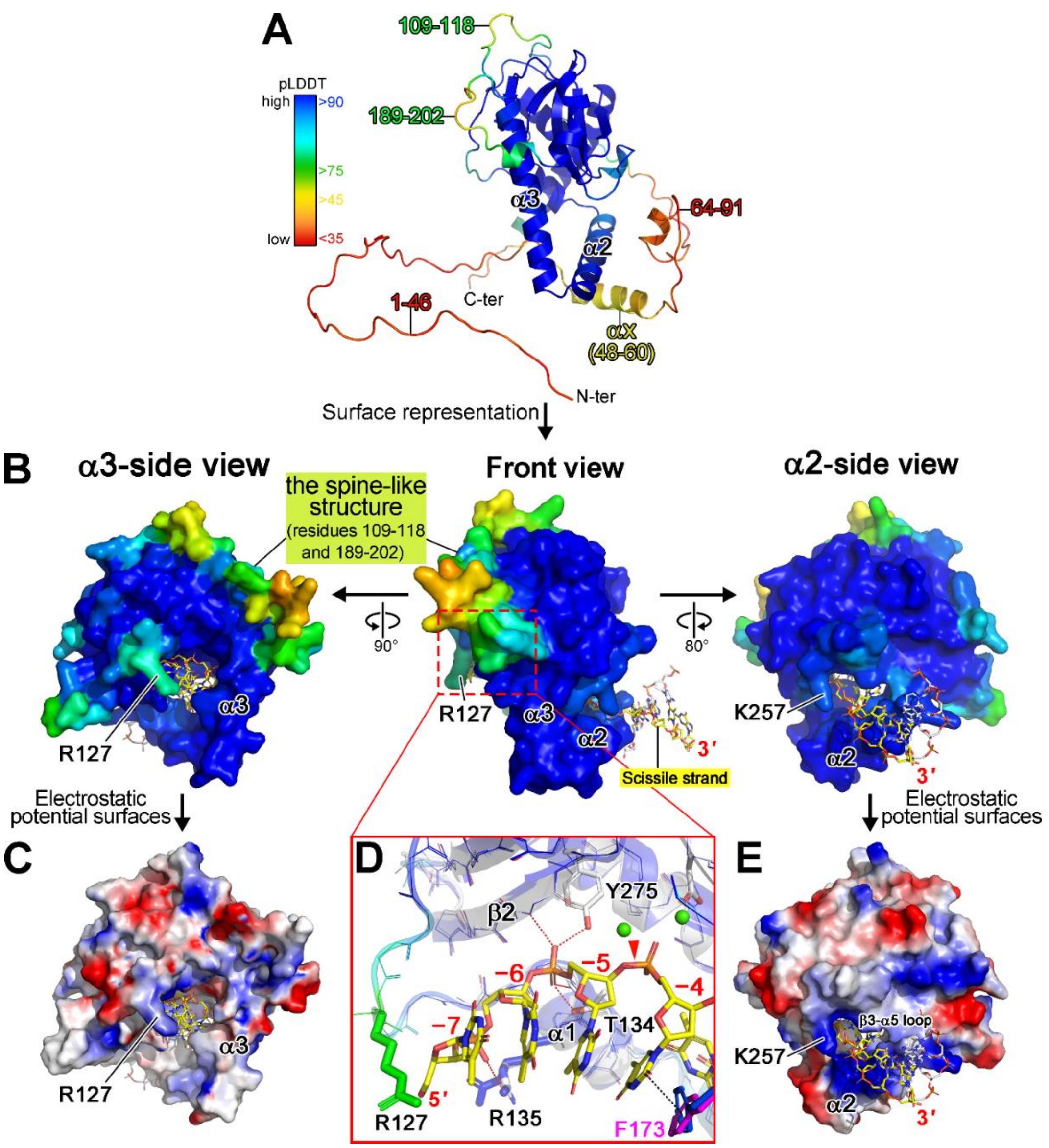


**Supplementary Figure S6.** Superimposition of the 5′-overhang DNA-bound MGME1 structure and AlphaFold2-predicted model. (**A**) The overall view of the AlphaFold2 (AF2)-predicted model of *Homo sapiens* MGME1 (4,5). The model is colored by per-residue confidence score (pLDDT) generated by AF2. (**B-E**) Superimposition of the 5′-overhang DNA-bound MGME1 structure and AF2-predicted model. The AF2 model was aligned with the protomer B of the 5′-overhang DNA-bound MGME1 structure as shown in Supplementary Figure S2B (RMSD=0.299 (over 181 $C_\alpha$)), and the protein of the latter structure is omitted, showing only the bound DNA (in stick representation) superimposed with the AF2 model. In panels **B-C**, the AF2 model is shown in surface representation and colored by pLDDT, thus highlighting the spine-like structure formed by the moderate pLDDT regions. In panels **C** and **E**, the electrostatic potential surfaces (generated in PyMOL) of the AF2 model are shown. Panel **D** is the enlarged view of the binding site of scissile strand nucleotides (nt) −4 to −7. In this view, both superimposed proteins are shown, with the AF2 model colored by pLDDT and the 5′-overhang DNA-bound structure colored in gray. The interactions between the protein and bound DNA backbone are illustrated by red dashed lines, while the black dashed line illustrates the stacking between F173 and the −4 nt. The $Ca^{2+}$ ions bound in the catalytic site are shown as green spheres. The red arrowhead indicates the scissile bond.

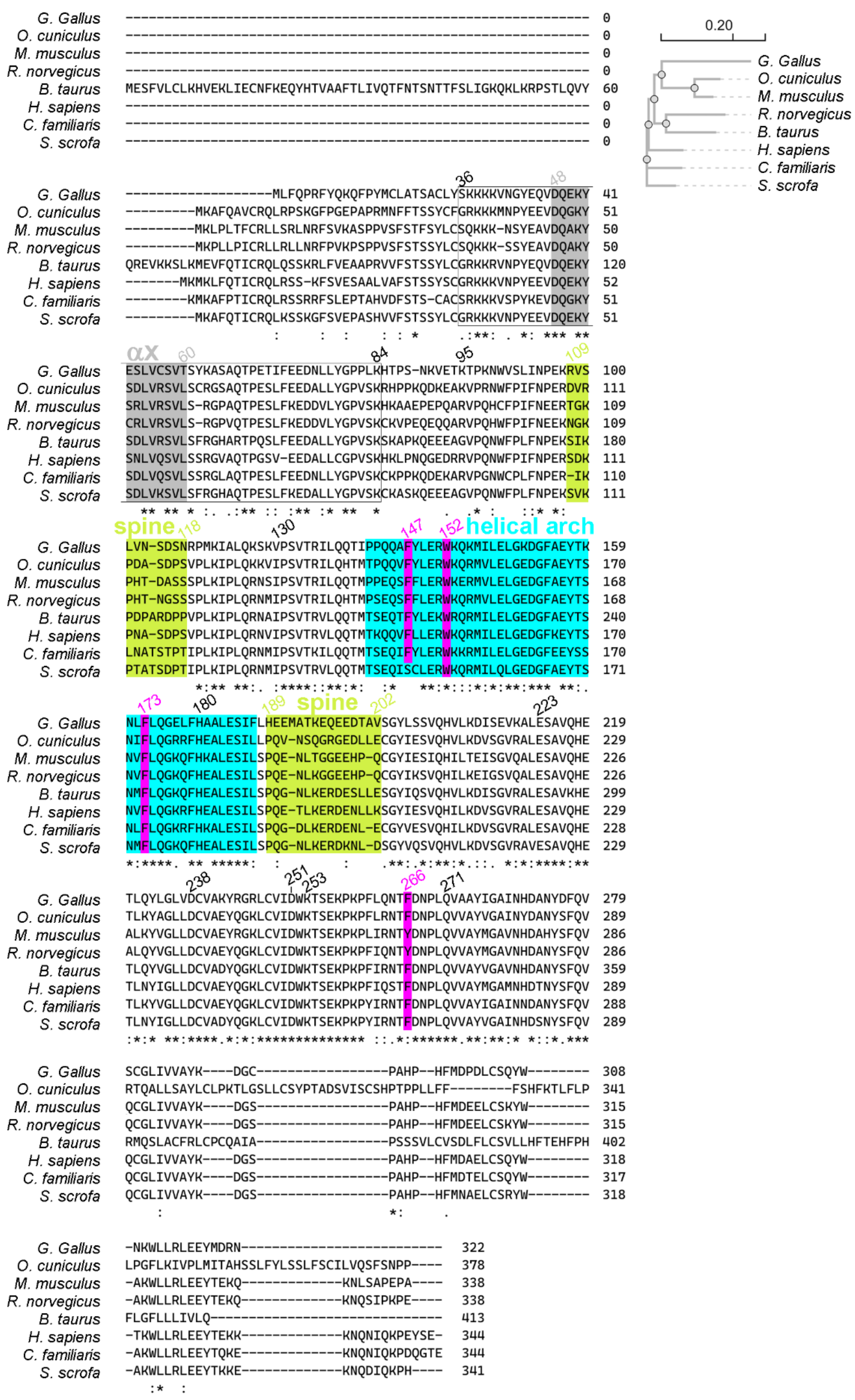


**Supplementary Figure S7.** Amino acid sequence alignment of MGME1 orthologs. The relatively conserved region (residues 36-84) in the dynamic N-terminus is enclosed by a box. The aligned MGME1 orthologs were acquired from *Gallus gallus* (*G. Gallus*), *Oryctolagus cuniculus* (*O. cuniculus*), *Mus musculus* (*M. musculus*), *Rattus norvegicus* (*R. norvegicus*), *Bos taurus* (*B. taurus*), *Homo sapiens* (*H.* sapiens), *Canis lupus familiaris* (*C. familiaris*), and *Sus scrofa* (*S. scrofa*).

### SUPPLEMENTARY REFERENCES


1. Wei, Z.H., Zhu, P. and Huang, Q.R. (2019) Investigation of ovotransferrin conformation and its complexation with sugar beet pectin. *Food Hydrocoll.*, **87**, 448-458.
2. Smilgies, D.M. and Folta-Stogniew, E. (2015) Molecular weight-gyration radius relation of globular proteins: a comparison of light scattering, small-angle X-ray scattering and structure-based data. *J. Appl. Crystallogr.*, **48**, 1604-1606.
3. The PyMOL Molecular Graphics System, V.S., LLC.
4. Jumper, J., Evans, R., Pritzel, A., Green, T., Figurnov, M., Ronneberger, O., Tunyasuvunakool, K., Bates, R., Zídek, A., Potapenko, A. *et al.* (2021) Highly accurate protein structure prediction with AlphaFold. *Nature*, **596**, 583-589.
5. Varadi, M., Anyango, S., Deshpande, M., Nair, S., Natassia, C., Yordanova, G., Yuan, D., Stroe, O., Wood, G., Laydon, A. *et al.* (2022) AlphaFold Protein Structure Database: massively expanding the structural coverage of protein-sequence space with high-accuracy models. *Nucleic Acids Res.*, **50**, D439-D444.
6. Tsutakawa, S.E., Thompson, M.J., Arvai, A.S., Neil, A.J., Shaw, S.J., Algasaier, S.I., Kim, J.C., Finger, L.D., Jardine, E., Gotham, V.J.B. *et al.* (2017) Phosphate steering by Flap Endonuclease 1 promotes 5′-flap specificity and incision to prevent genome instability. *Nat. Commun.*, **8**.
